# Architecture of the chikungunya virus replication organelle

**DOI:** 10.1101/2022.04.05.487153

**Authors:** Timothée Laurent, Pravin Kumar, Susanne Liese, Farnaz Zare, Mattias Jonasson, Andreas Carlson, Lars-Anders Carlson

**Affiliations:** Department of Medical Biochemistry and Biophysics, Umeå University, Umeå, Sweden; Wallenberg Centre for Molecular Medicine, Umeå University, Umeå, Sweden; Molecular Infection Medicine Sweden, Umeå University, Umeå, Sweden; Department of Mathematics, Mechanics Division, University of Oslo, N-0851 Oslo, Norway; Max Planck Institute for the Physics of Complex Systems, Dresden, Germany

**Keywords:** alphavirus, chikungunya, chikungunya virus, RNA, cryo-EM, cryo-electron tomography, plasma membrane, in vitro reconstitution, mathematical model

## Abstract

Alphaviruses are mosquito-borne viruses that cause serious disease in humans and other mammals. Along with its mosquito vector, the alphavirus chikungunya virus (CHIKV) has spread explosively in the last 20 years, and there is no approved treatment for chikungunya fever. On the plasma membrane of the infected cell, CHIKV generates dedicated organelles for viral RNA replication, so-called spherules. Whereas structures exist for several viral proteins that make up the spherule, the architecture of the full organelle is unknown. Here, we use cryo-electron tomography to image CHIKV spherules in their cellular context. This reveals that the viral protein nsP1 serves as a base for the assembly of a larger protein complex at the neck of the membrane bud. Biochemical assays show that the viral helicase-protease nsP2, while having no membrane affinity on its own, is recruited to membranes by nsP1. The tomograms further reveal that full-sized spherules contain a single copy of the viral genome in double-stranded form. Finally, we present a mathematical model that explains the membrane remodeling of the spherule in terms of the pressure exerted on the membrane by the polymerizing RNA, which provides a good agreement with the experimental data. The energy released by RNA polymerization is found to be sufficient to remodel the membrane to the characteristic spherule shape.

## Introduction

Chikungunya is a mosquito-borne disease characterized by a rapid onset of fever, followed by debilitating joint pains and arthritis that can last for months or years (1, 2). It is severely underdiagnosed, but suspected cases have surpassed 500,000/year in several recent years (https://www.who.int/news-room/fact-sheets/detail/chikungunya). The causative agent of chikungunya is chikungunya virus (CHIKV), a positive-sense single-stranded RNA virus of the alphavirus genus (family Togaviridae). In the last two decades, CHIKV has spread rapidly, far beyond its probable origins in east Africa, to cause large outbreaks in Asia and the Americas. One reason for this is its adaptation to a new mosquito host, *Aedes albopictus*, which inhabits more temperate regions (3, 4). In addition to CHIKV, a plethora of pathogenic alphaviruses exist, and their utilization of different mosquito species highlights the potential for new variants to arise and spread. There are no approved vaccines or antivirals against any alphavirus-caused diseases.

The replication of the alphavirus genome takes place in a virus-induced RNA replication organelle, also known as a “spherule” or “replication complex”. This organelle is formed as an outward-facing plasma membrane bud with a diameter of 50-80 nm (5). The size of the membrane bud has been shown to depend on the length of the viral genome (6). The bud is thought to have a stable, open neck that connects it to the cytoplasm, and this high-curvature membrane shape persists for several hours in the infected cell during active RNA production. The viral nsPs are thus thought to serve the additional role of maintaining this peculiar membrane shape while replicating the viral RNA.

The alphavirus genome codes for four non-structural proteins (nsP1-nsP4), initially produced as one polyprotein, with distinct functions in the viral genome replication (5, 7). NsP1 caps the 5’ end of the new viral RNA independently of the host-cell capping machinery (5). It is the only nsP reported to bind membranes, and its membrane affinity is enhanced by, but not dependent on, a palmitoylation site (8). NsP2 has RNA helicase and RNA triphosphatase activity in its N-terminal domain, and its C-terminus harbors a cysteine protease domain which cleaves the polyprotein into individual nsPs. NsP3 has ADP-ribosyl hydrolase activity, and interacts with several host-cell proteins (9). NsP4 is the RNA-dependent RNA polymerase directly responsible for the production of new viral RNA.

Structures have been determined for individual domains of the nsPs (10-12). Although informative for the function of the individual proteins, the structures generally provide no clues as to how the nsPs spatially coordinate the different steps of the RNA production and the membrane remodeling. One exception is the structure of the isolated, nsP1 protein (13, 14). When overexpressed in eukaryotic systems and gently extracted from the plasma membrane, nsP1 was shown to form a ring-shaped dodecamer, displaying its active sites to the inside of the ring and the membrane-binding surfaces to the outside. It was thus suggested that the nsP1 dodecamer may bind at and stabilize the high-curvature membrane neck. This model remains to be tested experimentally, and it is not known how localization of nsP1 at the neck would relate to other protein components in the spherule, the RNA or the membrane shape.

Here we use cellular cryo-electron tomography, *in vitro* reconstitution and mathematical modelling to provide a first integrated model of the CHIKV spherule. Our findings reveal that nsP1 anchors a large protein complex at the membrane neck, and directly recruits nsP2 to the membrane. The lumen of full-sized spherules contains a single copy of the viral genome, and we present a theoretical model that explains how RNA polymerization leads to a membrane remodeling consistent with the shapes observed in the tomograms.

## Results

### Cryo-electron tomography allows visualization of CHIKV spherules at the plasma membrane

We wished to study the structure of the CHIKV spherule *in situ* in unperturbed cells. The high biosafety level necessitated by CHIKV is typically dealt with by chemical fixation of infected cells prior to electron microscopy. Since this may compromise macromolecular organization, we instead opted to use viral replicon particles (VRPs), which transduce cells with a replication-competent, but capsid protein-deleted, genome that results in a self-limiting single-cycle infection (15). The VRPs express an eGFP reporter gene in place of the capsid proteins, which allowed confirmation that a vast majority of the cells grown on EM grids were transduced and had active viral RNA replication (Fig. S1). Cryo-electron tomograms of the peripheral plasma membrane occasionally showed CHIKV spherules appearing as clusters of balloon-shaped organelles sitting at the plasma membrane (Fig. 1A-B, Movie S1). They had a diameter ranging from 50 to 70 nm, consistent with what has been reported from resin-section EM (16). In addition to the membrane topology, the cryo-electron tomograms also revealed filamentous densities coiled on the inside of the membrane buds (Fig. 1A-B, Movie S1). The position and width of the filaments make is likely that they are the viral RNA, possibly in its dsRNA replicative intermediate. We next turned to the stabilization of membrane curvature. In principle, the high-curvature membrane of the CHIKV spherule could be stabilized either by protein binding throughout the curved membrane, or by specific stabilization of the membrane neck. From visual inspection, there was no consistent pattern of protein coating over the curved surface of the membrane bud. On the other hand, in all imaged spherules we observed a macromolecular complex sitting at the membrane neck (Fig. 1A-B). In well-resolved individual spherules, the complex seemed to be bipartite with a base pinching the neck of the spherule and a second part protruding toward the cytoplasm of the cell (Fig. 1C). Taken together, these data suggest that the CHIKV spherule consists of a membrane bud filled with viral RNA, and has a macromolecular complex gating the opening of this bud to the cytoplasm (Fig. 1D).

**Figure 1:**
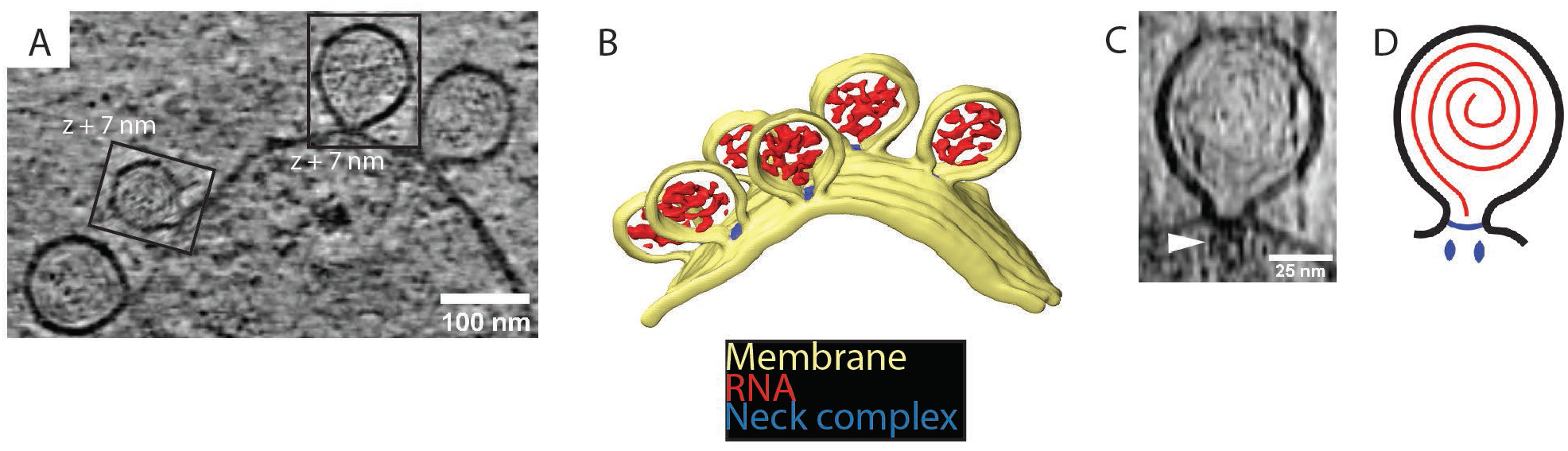
Cryo-electron tomography visualizes CHIKV spherules at the plasma membrane. (A) Computational slice through a cryo-electron tomogram of a CHIKV VRP-transduced BHK cell. The two framed insets are offset in the tomogram volume by 7 nm. (B) 3D segmentation of the tomogram shown in (A). Yellow: plasma membrane, Red: viral RNA, Blue: protein complex sitting at the spherule necks. (C) Subtomogram containing one spherule. The arrow indicates the densities present at the membrane neck. (D) Schematic of an initial model of the organization of a spherule. Black: plasma membrane, Red: viral RNA, Blue: protein complex sitting at the spherule necks.

### Subtomogram averaging determines the position of nsP1 in a larger neck complex

We were interested in investigating the structure of the protein complex sitting at the membrane neck. A 34 Å subtomogram average was calculated (Fig. S2) from 64 spherules without imposing any symmetry. It revealed that the complex is composed of two parts: a membrane-bound “base”, and a barrel-like “crown” (Fig. 2A-D). The base fits the membrane neck snugly (Fig. 2A-B). The crown is composed of three rings and protrudes towards the cytoplasm (Fig. 2A-C). At the current resolution there is no visible connection between the base and the crown. A third component of the neck complex is a central density protruding from the base, through the crown towards the cytoplasm. It appears more diffuse than the base and the crown.

**Figure 2:**
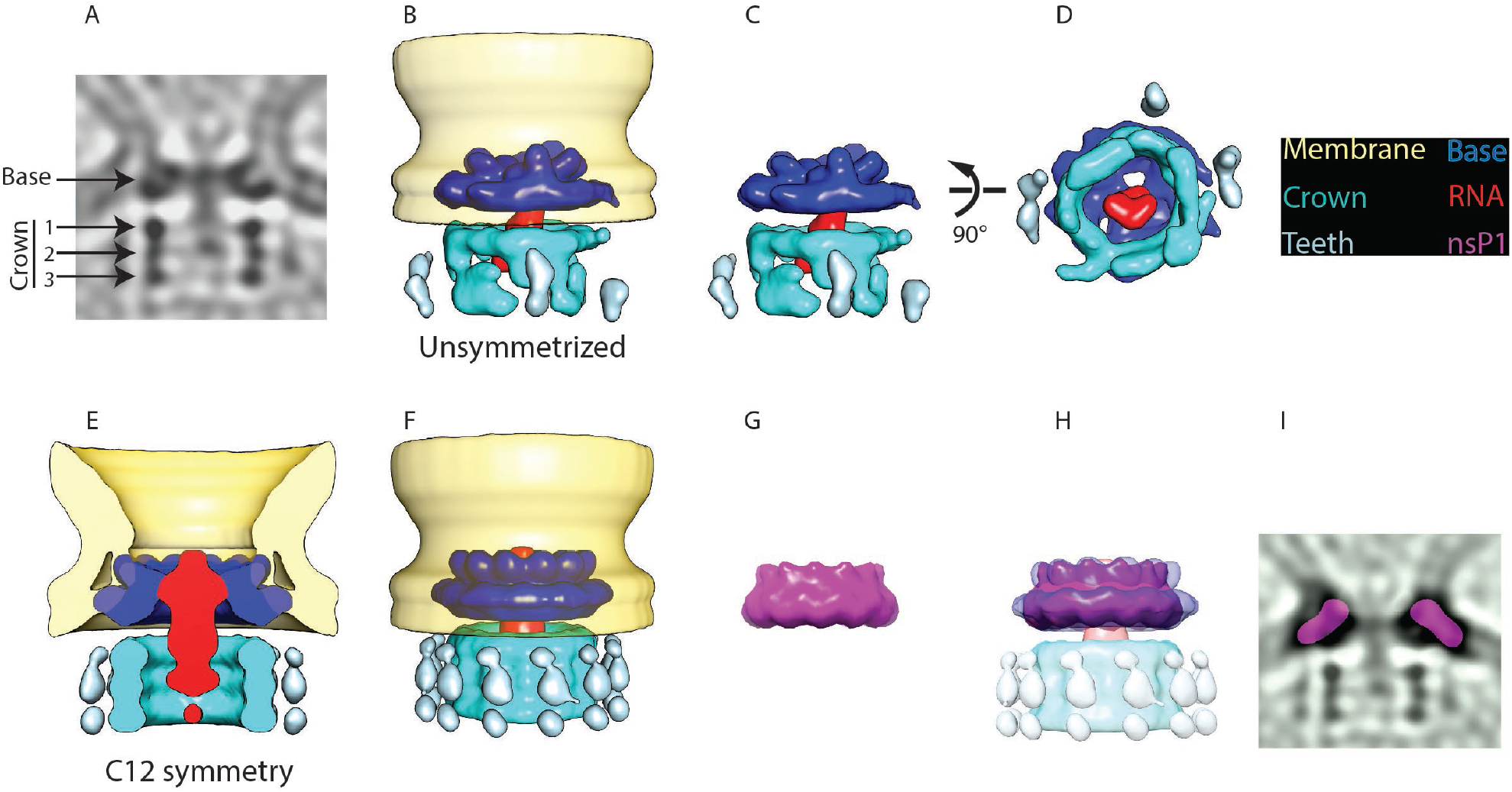
Subtomogram averaging reveals the multipartite nature of the neck complex. (A) Central slice through the unsymmetrized subtomogram average of the neck complex, low-pass filtered to its Gold-standard resolution 34Å. Arrows indicate densities that are referred to as the “base” and the “crown”. The crown is located on the cytoplasmic side and is composed of three rings. Density is black. (B) 3D segmentation of the unsymmetrized subtomogram average shown in (A). The spherule membrane (yellow) is radially symmetrized for clarity. Dark blue: base, red: putative RNA, cyan: crown, light blue: teeth. (C), (D): views of the neck complex in different orientations. (E) Color key for neck complex components. (F) Cross-section through the subtomogram average of the neck complex with C12 symmetry imposed, low-pass filtered to its Gold-standard resolution 28 Å. (G) Surface view corresponding to (F). (H) The structure of the isolated nsP1 (from (13)) low-pass filtered to the resolution of our average. (I) Superimposition of nsP1 onto the base of the protein neck complex. (J) Slice through the unsymmetrized subtomogram average, as in (A), with a slice of the fitted nsP1 superimposed on the base of the complex.

We hypothesized that the base of the neck complex may be nsP1, the only nsP with known membrane-binding motifs. The recent structures of nsP1 revealed a ring-shaped dodecamer with a similar dimension to the base of the neck complex (13, 14). For comparison, we imposed 12-fold symmetry on our neck complex (Fig. 2E-F) and low-pass filtered the published nsP1 structure to the 28 Å resolution of the 12-fold symmetrized average (Fig. 2H). An overlay of these two showed a close match in size and shape of the isolated nsP1 and the base of the neck complex (Fig. 2H). The best fit of nsP1 into the neck complex is such that the narrow side of the nsP1 ring, carrying the membrane-association sites, is in direct contact with the membrane. We further verified that nsP1 fits the unsymmetrized neck complex average (Fig. 2I). This overlay indicated that there may be additional densities bound to the inside of the nsP1 ring in the full neck complex as compared to the heterologously expressed nsP1.

There was not sufficient signal in the subtomogram average to experimentally determine the rotational symmetry in the crown part of the neck complex. But the main features were consistent between the unsymmetrized and the C12 averages: the crown consists of three stacked rings of equal diameter (Fig. 2A,E) and there is weaker but consistent density for peripheral structures (“teeth”) surrounding the rings (Fig. 2B,C,F). At the current resolution, the components forming the crown and teeth cannot be identified from the subtomogram average. However, based on their volume of 1500 - 1700 nm^3^ we estimate them to have a molecular mass of 1.2-1.4 MDa. At the center of the neck complex, extending out from nsP1 through the crown, is a rod-like density that is the only candidate to be the new viral RNA leaving the spherule. In summary, the subtomogram average suggests that nsP1 forms the base of a larger neck complex that extends towards the cytoplasm with a barrel-like structure that may funnel new viral RNA out from the spherule lumen.

### NsP1 recruits nsP2 to membranes containing monovalent anionic lipids

To establish the biochemical basis of the neck complex assembly through nsP1 we purified recombinant CHIKV nsP1 to homogeneity (Fig. S3). To test whether a monomeric nsP1 can bind the membrane prior to oligomerization, we used the monomeric fraction of nsP1 and synthetic liposomes in a multilamellar vesicle (MLV) pulldown assay (Fig S3). In the absence of any negatively charged lipids, nsP1 did not bind appreciably to the vesicles (Fig. 3A). Semliki forest virus nsP1 has been reported to associate with phosphatidyl serine (PS), an abundant lipid on the inner leaflet of the plasma membrane (17). Thus, we next decided to include PS in the MLVs. This revealed that nsP1 has concentration-dependent binding to PS-containing membranes (Fig. 3A, Fig. S4). The pulldowns were repeated in the presence of other monovalent anionic glycerophospholipids (phosphatidyl glycerol (PG) and phosphatidyl inositol (PI)), to test whether the binding was specific to PS or more generally dependent on membrane charge. NsP1 showed very similar, concentration-dependent binding to PG and PI-containing membranes (Fig. 3A). We then studied the interaction of nsP1 with phosphoinositides (PIPs), lipids that serve as membrane identity markers and may thus be involved in targeting the spherule assembly to a specific membrane. We compared two phosphoinositides: the predominantly Golgi-resident PI(4)P, and the predominantly plasma membrane-resident PI(4,5)P_2_. Curiously, nsP1 had a higher affinity to membranes containing low PIP concentrations, and almost no affinity for membranes with higher concentration of these lipids (Fig. 3B-C). For each PIP, we observed weaker membrane association than to membranes containing monovalent anionic lipids (Fig. 3A-C). As an alternative approach we visualized the interaction of nsP1 with giant unilamellar vesicles (GUVs) using confocal microscopy. No accumulation of fluorescent nsP1 was seen on GUVs consisting of phosphatidyl choline and cholesterol (a net-uncharged membrane). On the other hand, the equivalent charge density introduced in the form of either 20% PS or 5% PI(4,5)P_2_ led to visible binding of nsP1 to the surface of GUVs (Fig. 3D). 20% of the PI(4,5)P_2_-containing GUVs were positive for nsP1 binding, whereas 50% of PS-containing GUVs were positive, paralleling the MLV pulldown results (Fig. 3E).

**Figure 3.**
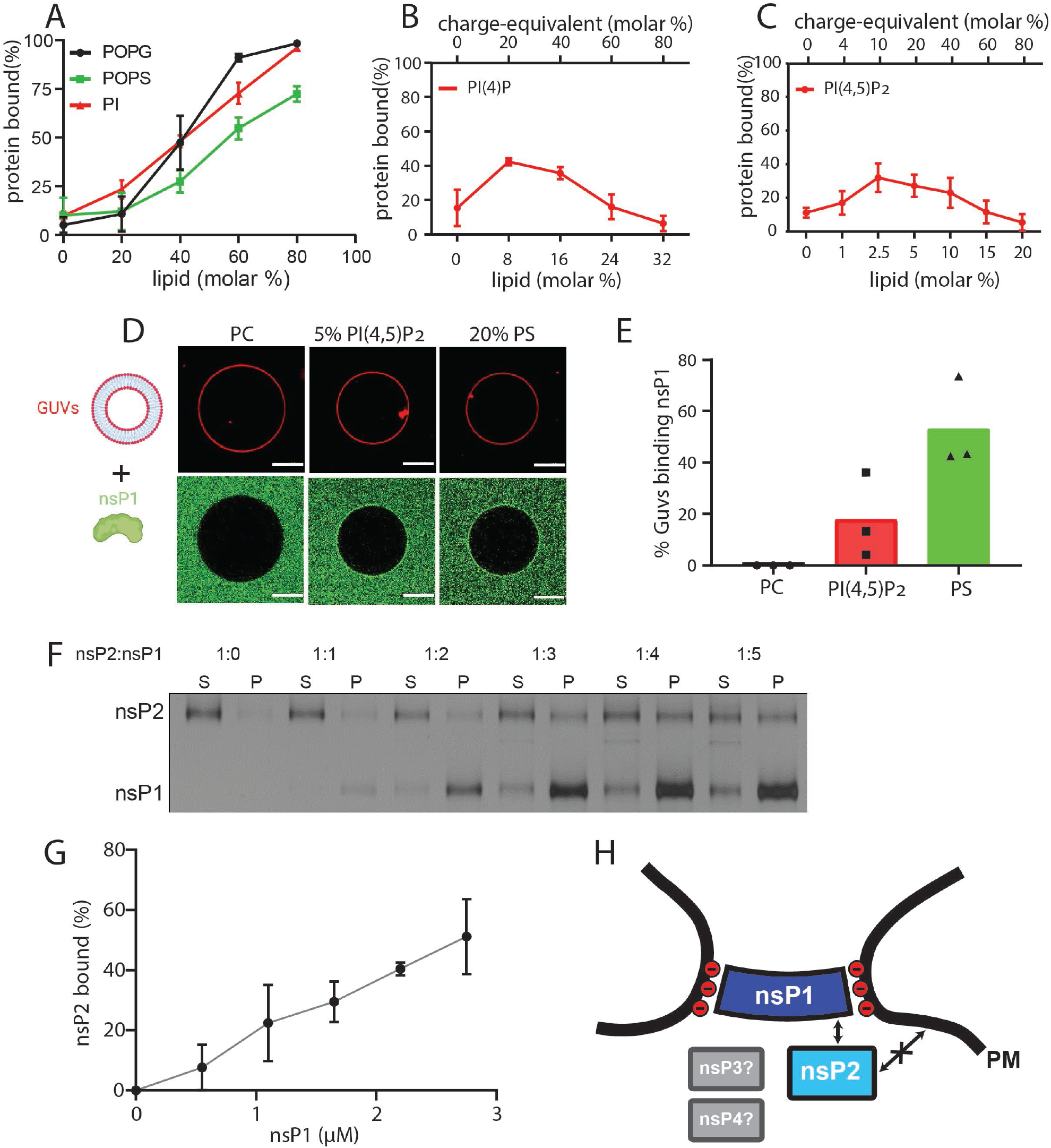
NsP1 binds to membranes containing monovalent anionic lipids and recruits nsP2 in a concentration-dependent manner. (A-C) Copelletation of nsP1 with multilamellar vesicles (MLVs) with varying percentages of the anionic phospholipids POPS, POPG, PI (A), PI(4)P (B), and PI(4,5)P_2_ (C) in a background of POPC and 20% cholesterol. Representative example gels shown in Fig. S4. The percentage of protein associated with membranes was quantitated from gels and plotted. Each plot represents the mean ± S.D of three independent replicates. (D) Confocal imaging of nsP1-ATTO488 (green) binding to GUVs (red) with POPC, or POPC including 5 mol % PI(4,5)P_2_, or 20% POPS. (E) Quantification of nsP1-bound GUVs from three experiment series. Data represents the percentage of nsP1-binding GUVs calculated from total number of GUVs observed for each experiment series plotted against the respective GUVs types. (F) Co-pelletation assay of nsP2 and nsP1 with POPS-containing MLVs. NsP2 and MLV concentrations were kept constant, while the nsP1 concentration was varied. Analysis of supernatant (S) and pellet (P) fractions by SDS-PAGE. G) Quantification of pelleted nsP2 with nsP1. The experiment shown in (F) was repeated two times. The pellet intensity at each nsP1 concentration was normalized to the total nsP2 intensity and plotted (mean ± S.D) against the nsP1 concentration. H) Schematic of the findings from (A-G), in the context of the neck complex. The non-structural proteins nsP3 and nsP4 were not included in these experiments but are displayed for completion as possible components of the neck complex.

The MLV pulldown assay was then extended to investigate if nsP1 can anchor other nsPs to the membrane. Both nsP3 and nsP4 have long disordered regions which make it challenging to obtain high-quality monodisperse protein. However, we were able to purify recombinant full-length nsP2 to homogeneity and obtained a monomeric protein (Fig. S3). In the pulldown assay nsP2 had no affinity to membranes containing 70 % PS. However, nsP2 was recruited to the membrane by nsP1 in a concentration-dependent manner (Fig. 3F-G). Taken together, these data show that the recruitment of nsP1 to membranes dependent mainly on monovalent anionic lipids, and that nsP1 can serve as a docking place for nsP2 that has no inherent membrane affinity (Fig. 3H).

### Full-size spherules contain a single copy of the genomic in dsRNA form

Turning next to the RNA component of the spherule, we reasoned that the visible filaments in the spherule lumen would allow an estimation of the total copy number of viral RNA within single spherules. The filaments were frequently observed to be relatively straight over a large fraction of the spherule lumen, which is more compatible with the persistence length of dsRNA (63 nm) than that of ssRNA (1 nm) (Fig. 1A-B, Movie S1) (18, 19). Using an automated filament tracing algorithm, we were able to trace long continuous stretches of dsRNA in the spherule lumen (Fig. 4A, Fig. S5). The traced model agreed well with filamentous densities seen in the tomograms, and the total filament length was robust over a wide range of parameter values (Movie S2, Fig. S6). We thus concluded that the filament tracing can be used to estimate the amount of genetic material present in a single spherule. Two tomograms of sufficiently high quality, recorded on different cells and each containing a cluster of full-sized spherules (n_1_=15 and n_2_=6), were analyzed. The total length of filaments for each data set were 18600±2900 Å/spherule and 21400±1600 Å/spherule (Fig. 4B). Assuming that the RNA was double-stranded and adopted an A conformation, the distance between two base pairs in this conformation is 2.56 Å (20, 21). Based on that assumption, the filament length corresponds to 7300±1150 bp and 8400±600 bp in the two tomograms, respectively (Fig. 4C). Since the length of the genomic RNA of our construct is 8820 bases, we thus conclude that there is on average nearly one copy of dsRNA within single, full-size spherules.

**Figure 4:**
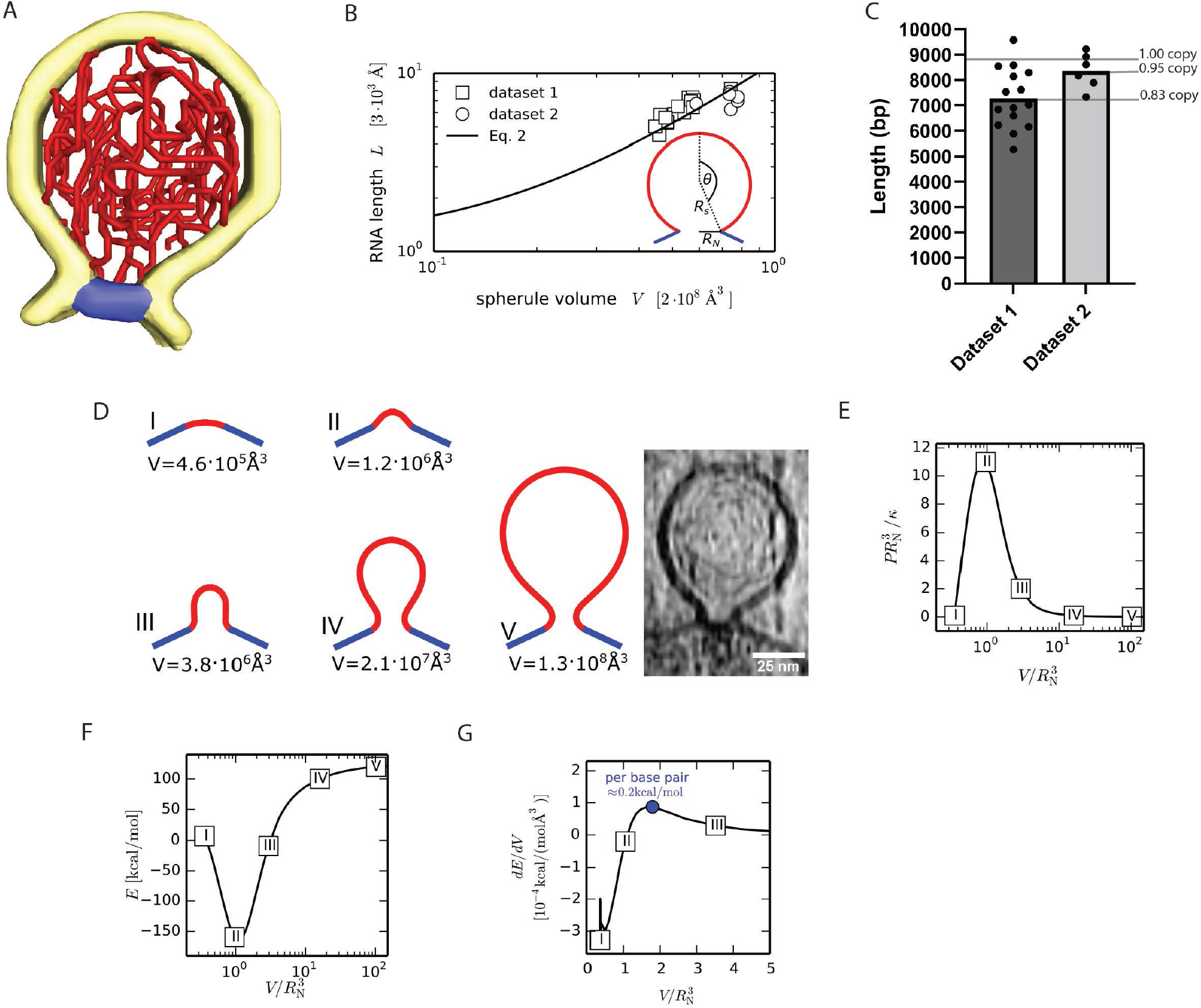
A single copy of the genomic RNA determines the shape of the spherule membrane. (A) Segmentation of the dsRNA traced within a spherule. Yellow: membrane, red: RNA, blue: neck complex base. (B) The RNA length L increases with spherule volume V. A common fit of both datasets with Eq. 2 gives *L*_0_ = (3 ± 1) ⋅ 10^3^Å and 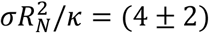⋅ 10^−2^, while *R*_*N*_ = 96 Å was determined experimentally. The inset shows the spherical cap model schematically. (C) Estimation of the dsRNA length (in base pairs) and the average copy number per spherule. One point represents a single spherule, and the datasets represents tomograms acquired on different cells. (D) Five shapes that minimize the energy (Eq. 1) for a given spherule volume. The last shave (V) is compared at scale with an experimental spherule tomogram. (E) Pressure-volume relation for a unitless membrane tension of a 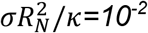. The corresponding membrane shapes are shown in subfigure D. (F) Energy (Eq. 1) as a function of the spherule volume for 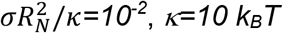 and *R*_*N*_ = 96 Å. (G) The energy change per change in volume is shown, which leads to a maximal energy to be supplied per base pair of 0.2 kcal/mol, where we assumed the volume of a single base pair to be not larger than 2 ⋅ 10^3^ Å^3^.

### The force exerted by RNA polymerization can drive spherule membrane remodeling

Proteins are known to induce membrane budding when they form spherical scaffolds that control the membrane shape (22-24). Since we observed viral proteins only at the spherule neck, we reasoned that other mechanisms may be involved in generating the characteristic high-curvature spherule membrane bud. To test this we developed a mathematical model of spherule membrane shape. The biological *Ansatz* of the model is that membrane remodeling is driven by the initial generation of dsRNA. This is the process in which the incoming positive-strand RNA is turned into a negative-strand copy which will be present in a duplex with the positive strand. This may happen in two ways: (i) the initial positive strand is present in a nascent spherule which grows as the single strand is turned into dsRNA, or (ii) the initial positive strand is translocated into the spherule lumen concomitant with the production of the complementary negative strand. Either of (i) and (ii) are compatible with the model described below. The physical assumptions of the model are the following: We describe the membrane as a thin elastic sheet in a Helfrich-type model. The RNA, which is modeled as a semiflexible polymer, exerts a pressure onto the membrane that causes the spherule to expand.

As the dsRNA is produced it exerts a pressure *P* on the spherule membrane. The pressure that acts to increase spherule volume is balanced by the elastic membrane properties. To model the formation of a spherule, we begin by formulating the membrane energy *E*, which includes the Helfrich bending energy, the membrane tension *σ*, and the pressure *P* exerted by the viral RNA (25).

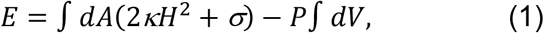

with the membrane area *A*, the bending rigidity *κ*, the mean curvature *H* and the spherule volume *V*.

To derive a scaling relation between the RNA length *L* and the spherule volume *V*, we approximate the shape by a spherical cap, as indicated in the inset of Fig. 4B, where the radius *R*_*s*_ and the polar angle *θ* are related via the neck radius *R*_*N*_*=R*_*s*_*sin(θ)*. In the limit of large spherule (*θ≈π*) we find 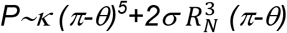 and *V∼(π-θ)*^*-3*^ (see Supporting Information). It is known from polymer theory that the pressure volume relation of long semiflexible polymers in spherical confinement scales to leading order as *PV∼LV*^*-2/3*^ (26-28). Hence, the RNA length scales with the spherule volume as a power law with

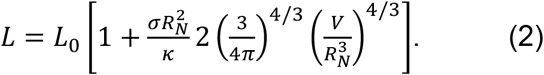

The neck radius is determined from EM imaging with *R*_*N*_ = 96 Å. Based on the data shown in Fig. 4B) we fit a value of *L*_0_ = (3 ± 1) ⋅ 10^3^ Å for the prefactor in Eq. 2 and a scaled membrane tension 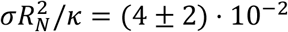. For comparison, with a bending rigidity of *κ=10 k*_*B*_*T*, we obtain *σ=10*^*-5*^ *N/m*, within the range of experimentally measured membrane tensions (29, 30).

Next, we study the membrane shape transformation from an initial pit to a fully formed spherule. The energy (Eq. 1) is minimized using the Euler-Lagrange method (see Supporting Information). To this end, we apply the arc-length parameterization and constrain the membrane in the neck region to the experimental geometry of the neck complex. In Fig. 4E the pressure-volume-relation is shown, for 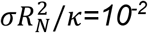. We find that the largest pressure is exerted for a rather small membrane pit with a volume of 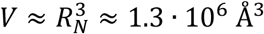. With a bending rigidity of *κ=10 k*_*B*_*T*, we find by solving Eq. 1 that an energy barrier of roughly 250 kcal/mol has to be overcome going form a flat membrane to a fully formed spherule (Fig. 4F). However, the energy cost per RNA base pair is much smaller. In Fig. 4G the change in energy per change in volume in shown. We see a maximum of 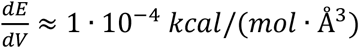 around 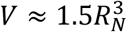. Assuming that each additional RNA base pair increases the volume by at most 2 ⋅ 10^3^ Å^3^, we estimate the maximum energy to be supplied per base pair at 0.2 *kcal/mol* at 25°C. The free energy change of RNA polymerization, including hydrogen bonding with the template, amounts to *ΔG*_0_ = −1.9 *kcal*/(*mol* ⋅ *base*) without accounting for the hydrolysis of the pyrophosphate. Comparing the two we conclude that the free energy released by RNA polymerization is around ten times larger than the energy required to bend the membrane, even at its peak “resistance”. Thus, RNA polymerization is sufficient to remodel the spherule membrane into its characteristic shape, assuming the neck geometry is constrained.

## Discussion

In this study we investigated the structural organization of spherules, the RNA replication organelles of alphaviruses. Our main findings are summarized in figure 5. Four viral proteins, nsP1-nsP4, are involved in the alphavirus genome replication (5). High-resolution structures have been determined for isolated domains of several nsPs, and for a ring-shaped dodecamer of the capping enzyme nsP1, the only nsP known to have membrane affinity (10, 11, 13, 14, 31, 32). These structures have provided insights into individual viral enzymatic functions, but not their cellular structural context, i.e. the spherules. Alphaviruses are not only a major source or morbidity, but their unique RNA replication mechanism is also used to develop self-replicating RNA vaccines that induce a more potent immune response than conventional mRNA vaccines (33). Underlying both the pathogenic viruses and the self-replicating RNA vaccine candidates is the same spherule machinery, which highlights the importance of understanding its organization. Our subtomogram average of the spherule neck complex (Fig. 2) provides first insights into this and suggests that the ring-shaped nsP1 dodecamer serves as the assembly hub for a larger protein complex sitting at the neck of the membrane bud. Complementary biochemical reconstitution showed that nsP1 is necessary and sufficient for membrane association of the helicase-protease nsP2.

**Figure 5:**
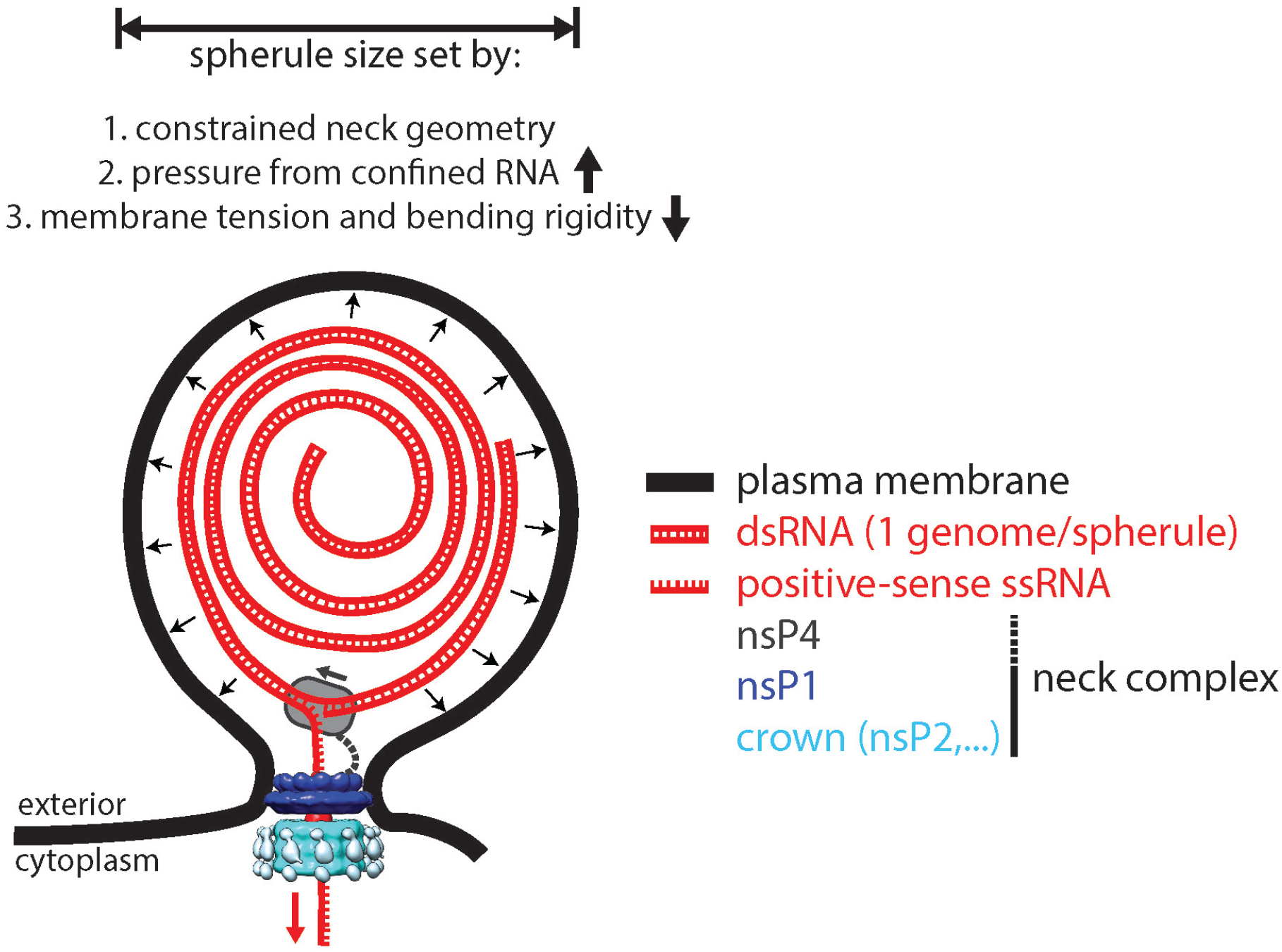
A model for the interplay between membrane, RNA and proteins in the CHIKV spherule. Each spherule contains a single viral genome, to >80% present in dsRNA form. The membrane shape is determined by the confined neck geometry, the pressure exerted by the confined genome, and the tension and stiffness of the membrane. NsP1 determines the neck geometry and serves an base plate for assembly of an additional 1.2 MDa complex. Biochemical evidence indicates that nsP2 is part of the neck complex, and an association of the viral polymerase nsP4 with the neck complex, although not directly addressed in this article, would be consistent with the suggested strand-displacement replication mode that produces several positive-sense strands from each spherule.

Our data align alphaviruses with an emerging theme in positive-sense RNA virus replication: macromolecular complexes located at a membrane neck play key roles in genome replication. All positive-sense RNA viruses utilize cytoplasmic membranes to compartmentalize their RNA replication machineries. It has been suggested to split the replication compartments into two groups based on membrane topology: double-membrane vesicles and membrane buds. This grouping based on membrane topology has recently been shown to correlate with the newly proposed phyla Pisuviricota and Kitrinoviricota, respectively (34). Alphavirus spherules fall into the latter category together with e.g. flaviviruses and nodaviruses. Whereas nothing is yet known about any flavivirus neck complex, cryo-electron tomography has revealed that nodaviruses have a neck complex of similar dimensions as the alphavirus neck complex we present here (35, 36). Strikingly, even the double-membrane vesicles-type replication organelles of coronaviruses have a neck complex connecting their lumen with the cytosol (37). The degree of structural conservation between neck complexes remains to be determined, but they have likely all evolved to solve the same problem: creating an environment conducive to viral genome replication in a cytoplasm rife with antiviral defense systems.

The limited resolution of the subtomogram average prevented us from determining if all enzymatic functions needed for RNA replication are localized directly in the neck complex. Conceptually, the localization of the polymerase nsP4 to the neck complex would be easy to reconcile with a strand-displacement mechanism that couples RNA polymerization to the extrusion of the displaced strand through the neck complex into the cytoplasm (Fig. 5). An eventual high-resolution structure of the entire neck complex may resolve this question, but a complete structural understanding of alphavirus RNA replication is likely to require several such structures due to the existence of a distinct early “negative strand” complex (38).

Analysis of the cryo-electron tomograms gave a clear answer to the question of the membrane bud contents: the lumen of full-size spherules consistently contains 0.8-0.9 copies of the viral genome in dsRNA form. While it has generally been speculated that the lumen of bud-type replication organelles contain the replicative intermediate, the consistency of the copy number is a striking outcome of our analysis. This may also suggests an explanation for why bud-type replication organelles all tend to have a similar diameter: they all contain genomes of ∼8-12 kb, which at a given density would all occupy a similar volume. Our mathematical model, consistent with the tomographic data, shows that the pressure exerted by the confined dsRNA, and the restriction of the neck geometry are sufficient to maintain the high-energy shape of the spherule membrane. Crucially, the model also show that the energy released by RNA polymerization is sufficient to drive the membrane shape remodeling. This establishes polymerase-driven budding as a new membrane remodeling mechanism. Future studies may determine if this mechanism is generally used in the large number of positive-sense RNA viruses with “bud-type” replication organelles. Taken together, our study takes the first steps towards an integrated structural model of an entire viral replication organelle, suggesting a high degree of spatial coordination of proteins, RNA and membrane components of the alphavirus spherule.

## Acknowledgements

We thank Andres Merits (Tartu) for sharing the CHIKV VRP system, and Tero Ahola (Helsinki) for insightful discussions at the early stage of this project. Cryo-EM and fluorescence microscopy were performed at the Umeå Centre for Electron Microscopy (UCEM) and Biochemical Imaging Center Umeå (BICU), respectively. UCEM is a SciLifeLab National Cryo-EM facility supported by instrumentation grants from the Knut and Alice Wallenberg Foundation and the Kempe Foundations. UCEM and BICU are part of the National Microscopy Infrastructure, NMI (VR-RFI 2016–00968). This project was funded by a Human Frontier Science Program Career Development Award (CDA00047/2017-C) to L.-A.C, the Knut and Alice Wallenberg Foundation (through the Wallenberg Centre for Molecular Medicine Umeå), the Swedish research council (grants 2018-05851, 2021-01145) and a Kempe foundations postdoctoral fellowship to P.K.

## Data Availability

The subtomogram averages of the neck complex have been deposited at the Electron Microscopy Data Band with accession codes EMD-XXXX (unsymmetrized) and EMD-XXXX (C12-symmetrized).

## Materials and Methods

### Cell culture

Baby Hamster Kidney (BHK) cells were cultured in Minimum Essential Medium (MEM, Gibco) supplemented with GlutaMAX (Gibco) and 10% Fetal bovine serium (FBS, Gibco). Cells were grown in an incubator at 37°C with 5% CO_2_.

### Viral replicon particles

The Viral Replicon Particles (VRPs) system consist of the viral genomic RNA depleted in structural proteins, and the helper plasmids C and E coding respectively for the capsid and the E3, E2, 6K, E1 structural proteins (15). The plasmids were kindly provided by Andres Merits, Tartu. The viral genomic RNA and the RNA of the helper plasmids C and E were *in vitro* transcribed and capped using the mMESSAGE mMACHINE SP6 Transcription kit (Thermo Fisher Scientific). Quality of the RNA was assessed on a denaturing formaldehyde-agarose gel. BHK cells were electroporated with the three RNA using the NEON electroporation system (Invitrogen). Cells were passaged one day prior to electroporation. Cells were then trypsinized and washed twice in PBS before being resuspended in the R resuspension buffer at a density of 10^7^ cells/mL and electroporated at 1200 V, 30 ms width, one pulse. Electroporated cells were then resuspended in antibiotic-free MEM medium supplemented with 10% FBS and transferred to a T75 flask. After 48h, the medium containing the VRPs was harvested, spun down to remove detached cells and cell debris. The VRP-containing supernatant, was aliquoted, flash-frozen on liquid nitrogen and stored at -80°C.

### Sample preparation

QUANTIFOIL R 2/1 Au 300 EM grids were glow discharged for 10 min at 15 mAh, sterilized and then set at the bottom of an IBIDI µ-Slide 8 well. Cells were seeded at 25,000 cells/well and left overnight to attach and spread on the EM grids. Cells were then transduced by swapping the cell medium for 250 µL of the VRP suspension. 6 h later, a solution of 5 nm Protein A-coupled colloidal gold (CMC-Utrecht) was added to each grid after which it was immediately plunge frozen in liquid propane-ethane using a FEI Vitrobot.

### Cryo-electron tomography

Data collection parameters are summarized in table S1. Vitrified cells were imaged using a transmission electron microscope, the FEI Titan Krios with an accelerating voltage of 300kV, a Gatan Bioquantum LS energy filter, a K2 summit detector. Tiltseries were acquired using SerialEM software (39), at a magnification of 33,000x in with a super-resolution pixel size of 2.19 Å/px. Data were gathered at the plasma membrane of infected BHK cell using either a bilateral or a dose-symmetric scheme (40) at a defocus between -3 and -5 µm. Typically, the total electron dose on the specimen was between 80 and 120 electrons/Å^2^, samples were tilted between -60° and 60° with an increment of 2°.

### Tomogram reconstruction

Movies generated during the data acquisition were motion corrected using MotionCor2 (41). Tiltseries were aligned using IMOD (42) based on 5 nm gold fiducials present on the specimen. The CTF was estimated using CTFFIND4 (43) and corrected using IMOD’s ctfphaseflip. The images were dose filtered (44) and tomograms generated using weighted back projection in IMOD.

### Subtomogram averaging

The subtomogram averaging was carried out as schematically indicated in Fig. S2. 76 particles were extracted from nine high-quality unbinned tomograms using Dynamo (45, 46). Of these 76 particles, 64 could be unambiguously oriented and centered manually before generating a first average of the protein neck complex. A cylindrical mask centered on the protein neck complex was created and a second round of alignment was performed allowing for full azimuthal rotations and limited (+/-30°) tilts with respect to the z axis (defined as the axis passing through the neck complex). Azimuthal angles of the particles in the crop table were then randomized in order to decrease the impact of the missing wedge and, by this process, another average was generated. This average was then used in combination with the original particle poses and a tighter cylindrical mask to obtain a third average. A custom mask was then defined on the center slice of the third average, radially symmetrized and used in a final alignment, still allowing full azimuthal rotations and limiting tilts and shifts. The final alignment was performed separately, once without symmetry and once with 12-fold rotational symmetry imposed. The resolutions were estimated to 34 Å and 28 Å for the unsymmetrized and C12-symmetrized averages, respectively, using the Gold-standard Fourier Shell Correlation with a threshold of 0.143.

### Creation of the segmented 3D models

The segmentation in Fig. 1B was created by manual segmentation in Amira (Thermo Fisher Scientific). For the subtomogram average of the neck complex, symmetrized and non-symmetrized averages were first filtered to their respective resolution and the tight mask was applied to them. A representation of the membrane neck was generated by applying C36 symmetry to the average, masking away the neck complex and then applying a Gaussian filter. Both symmetrized and non-symmetrized averages were segmented in UCSF Chimera (47) and the membrane template and averages were superimposed. The published structure of nsP1 (pdb 6z0v, reference (13)) was filtered to the resolution of the average and then fit in the density of the base of the neck complex using UCSF Chimera.

### Molecular mass estimation of crown subcomplex

The crown subcomplex was cropped out of the protein neck complex using the volume eraser function of Chimera. The volume of the cropped density was computed and the molecular weight was estimated assuming 825 Da/nm^3^ (48).

### Filament tracing

Binned tomograms were filtered using a SIRT-like filter with two iterations in IMOD and were imported in Amira where the RNA tracing was performed using its filament tracing functionality (49). Single spherules were cropped from the imported tomograms and a non-local means filter was applied applied to the cropped subtomograms with parameters selected to yield a clear contrast between the filament contained in the spherules and the background. A cylinder correlation was run with the filament width chosen to match dsRNA. The interior of spherules was segmented in order to leave out spurious hits in membranes and the exterior. Correlation lines were then traced with parameters selected to yield a good match between traces and visible filaments. The total filament length (in Å) as stated by the software was used to calculate dsRNA length in basepairs, assuming 2.56 Å/bp.

### Plasmids for protein production

Plasmids for CHIKV nsp1 and nsP2 were obtained by cloning codon-optimized CHIKV nsP1 and nsP2 genes of LR 2006_OPY1 strain into 2Bc-T vector (ORF-TEV-His6) and 1M vector (His6-MBP-TEV-ORF) respectively from Macrolab (University of California, Berkeley, USA).

### Expression and purification of CHIKV nsP1

To overexpress CHIKV nsP1, nsP1 plasmid was transformed into E. coli BL21(DE3) cells. An overnight culture was grown in Luria Broth (LB) supplemented with 100 μg/mL of carbenicillin at 37°C to inoculate the secondary culture. Cells were grown at 37°C to an O.D_600_ of 0.4 then the incubator temperature was reduced to 20°C. After the culture cooled down to 20°C and O.D_600_ reaches between 0.8-0.9, protein expression was induced with 0.5 mM Isopropyl β- d- 1-thiogalactopyranoside (IPTG) and continue the expression at 20°C overnight. Cells were harvested by centrifuging at 6000 rpm (rotor JLA-8.1000 Beckman Coulter, Brea, USA) for 60 min. After discarding the supernatant, cell pellet was washed with lysis buffer (50 mM Tris-HCl, pH 7.4, 500 mM NaCl, 0.1 mM THP, 36 μM NP40, 5 mM MgCl2, and 10% glycerol) and stored at – 80°C.

The entire purification of CHIKV nsP1 was performed at 4°C (either in the cold room or on ice). Cell pellets were thawed and resuspended in lysis buffer supplemented with DNase I and protease inhibitor cocktail (in-house preparation). Homogenized suspension then passed twice through a cell disruptor (Constant System Limited, Daventry, England) at a pressure 27 kPsi. Lysed cells was centrifugated at 21000 rpm (rotor JA-25.50 Beckman Coulter, Brea, USA) for 1 hour and the supernatant constituting the soluble fraction was passed through a 0.22 μm syringe filter to get a clear lysate. The cleared lysate was incubated for 2 h at 4°C on a rotating wheel with 1 ml Ni-Sepharose Fastflow resin (Cytiva) that was pre-equilibrated with lysis buffer. After incubation, lysate-resin suspension was loaded onto a 20 ml polypropylene gravity-flow column (Bio-Rad). After collecting the flow through, the protein-bound resin was washed with wash buffer (50 mM Tris-HCl, pH 7.4, 500 mM NaCl, 0.1 mM THP, 36 μM NP40, 5 mM MgCl_2_, 10 % glycerol, and 20 mM Imidazole) twice, each with 20 column volume (CV). Washed resin was resuspended in four-column volumes of lysis buffer and incubated after adding TEV protease (approx. 70 μg/ml; In-house prep) for overnight at 4°C on a rotator wheel. The cleaved protein was collected as flowthrough. An additional wash with 5 ml of lysis buffer was performed to collect the residual cleaved protein. Both elutions were pooled and further purified by Affinity chromatography. After diluting by adding buffer A (50 mM Tris-HCl, pH 7.4, 100 mM NaCl, 0.1 mM THP, 36 μM NP40, 5 mM MgCl_2_, and 10 % glycerol), diluted sample was filtered using 0.22 μM syringe filter (VWR) and loaded onto a HiTrap Heparin HP 1ml column (GE healthcare) pre-equilibrated with buffer A. Protein was eluted over a 14 CV NaCl gradient starting at 100 mM to a final 1M NaCl. Elutions were pooled down and concentrated using Vivaspin 6 centrifugal unit with 30 kDa cut off membrane (EMD Millipore) before being loaded onto a Superdex 200 increase 10/300 GL size-exclusion column (Cytiva) that was pre-equilibrated with SEC buffer (20 mM Tris-HCl, pH 7.4, 300 mM NaCl, 0.1 mM THP, and 5% glycerol). Protein elutions corresponding to monomeric-nsP1 peak were pooled and concentrated. Aliquots were then flash froze in liquid N_2_ and stored at -80°C.

### Expression and purification of CHIKV nsP2

Overexpression of CHIKV nsP2 was performed using LEX bioreactor in the following manner. The nsP2 plasmid was transformed into E. coli BL21(DE3) cells. An overnight culture was grown in Luria Broth (LB) supplemented with 50 μg/mL of kanamycin at 37°C to inoculate the secondary culture. Before going to the LEX Bioreactor, Terrific Broth (48.2 g per liter of TB supplemented with glycerol at 8 ml per liter) was augmented with Kanamycin (50 μg/mL) and antifoaming agent (approximately 15 drops in 1.5 L media; Sigma Aldrich). The media in 2 L bottles were kept at 37°C with bubbling for approx. 45 min and then inoculated with overnight primary culture (1:100). Around the O.D_600_ 0.35-0.45, changed the temperature of the bioreactor to 18°C and let the culture to cool down to 18°C. At this point, protein expression was induced with 0.5 mM IPTG and expression continued at 18°C for 18- 20 hrs. Cells were harvested by centrifuging at 6000 rpm (rotor JLA-8.1000 Beckman Coulter, Brea, USA) for 60 min. After discarding the supernatant, cell pellet was washed with lysis buffer (50 mM Tris-HCl, pH 8.0, 500 mM NaCl, 10 % glycerol, 0.1 mM THP, 36 μM NP-40) and stored at -80°C. The entire purification of CHIKV nsP2 was performed at 4°C. Cell mass was thawed and resuspended in lysis buffer supplemented with DNase I and protease inhibitor cocktail (in-house preparation). Homogenized suspension then passed twice through a cell disruptor (Constant System Limited, Daventry, England) at a pressure 27 kPsi. Lysed cells was centrifuged at 21000 rpm (rotor JA-25.50 Beckman Coulter, Brea, USA) for 1 hour and the supernatant constituting the soluble fraction was passed through a 0.22 μm syringe filter to get a clear lysate. The cleared lysate was incubated for 2 h at 4°C on a rotating wheel with 1 ml Talon Fastflow resin (Cytiva) that was pre-equilibrated with lysis buffer. After incubation, lysate-resin suspension was loaded onto a 20 ml polypropylene gravity-flow column (Bio-Rad). After collecting the flow through, the protein-bound resin was washed with wash buffer (50 mM Tris-HCl, pH 8.0, 500 mM NaCl, 0.1 mM THP, 36 μM NP40, 10 % glycerol, and 20 mM Imidazole) thrice, each with 20 column volume (CV). Protein was eluted with elution buffer (50 mM Tris-HCl, pH 8.0, 500 mM NaCl, 0.1 mM THP, 36 μM NP40, 10 % glycerol, and 250 mM Imidazole) in two fractions each of 5 ml. 6xHis-tag was removed by adding TEV protease (approx. 70 μg/ml; in-house prep) for overnight at 4°C on a rotator wheel. The cleavage mixture was centrifuged at 4000 rpm/4°C for 45 min to remove the visible precipitation. The supernatant was filtered using a 0.22 μM syringe filter and then pass through a HiTrap MBP-1 ml column pre-equilibrated with elution buffer to get rid of the His-MBP and His-TEV. The flowthrough, after diluting with buffer A (50 mM Tris-HCl, pH 8.0, 50 mM NaCl, 10 % glycerol, and 0.1 mM THP), was filtered using 0.22 μM syringe filter and loaded onto a HiTrap Heparin HP 1ml column (GE healthcare) pre-equilibrated with buffer A. Protein was eluted over a 14 CV NaCl gradient starting at 100 mM to a final 1M NaCl. Elutions were pooled down and concentrated using Vivaspin 6 centrifugal unit with 30 kDa cut off membrane (EMD Millipore) before being loaded onto a Superdex 200 increase 10/300 GL size-exclusion column (Cytiva) that was pre-equilibrated with SEC buffer (50 mM HEPES-NaOH, pH 8.0, 300 mM NaCl, 10% glycerol, and 0.1 mM THP). Protein elutions corresponding to nsP2 peak were pooled and concentrated. Aliquots were then flash froze in liquid N2 and stored at -80°C.

### Fluorophore labeling of CHIKV nsP1

Fluorophore labelling was performed on the eluent of the Heparin affinity chromatography. For labelling, the purification of nsP1 from metal-based affinity chromatography to Heparin affinity chromatography was performed in same buffers but Tris-HCl was replaced with HEPES-NaOH. The CHIKV nsP1 was mixed with 3-fold molar-excess of ATTO488 NHS (ATTO-TEK) and incubated at R.T. for 2 hrs. The free dye in the reaction was quenched by adding 1 M Tris-Cl, pH 7.4 to a final concentration of 50-100 mM and incubated further for 15-30 minutes. The CHIKV nsP1 labelling reaction was run through the HiLoad 16/600 Superdex 200 pg column pre-equilibrated with SEC buffer (20 mM Tris-HCl, pH 7.4, 300 mM NaCl, 0.1 mM THP, and 5% glycerol) to separate the monodisperse state of the labeled protein from the free dye. Labeling efficiencies were normally 70–100%.

### Liposome preparation

The phospholipids, for liposome preparation, 1-palmitoyl-2-oleoyl-sn-glycero-3-phospho-l-serine (POPS), 1-palmitoyl-2-oleoyl-sn-glycero-3-phosphocholine (POPC), 1-palmitoyl-2-oleoyl-sn-glycero-3-phospho-(1′-rac-glycerol) (POPG), L-α-phosphatidylinositol (PI)(Liver, Bovine), L-α-phosphatidylinositol-4-phosphate (PI(4)P)(Brain, Porcine), L-α-phosphatidylinositol-4,5-bisphosphate (PI(4,5)P_2_) (Brain, Porcine) were purchased as chloroform (or chloroform:methanol:water) solutions from Avanti Polar Lipids Inc. Cholesterol powder was purchased from Sigma and dissolved in chloroform.

### Multilamellar vesicles (MLVs)

MLVs were prepared by mixing phospholipids dissolved in solvent at the desired molar ratio (see Table S2). POPC was the bulk lipid, cholesterol was kept fixed at 20 mol %, and charged lipids were added to the desired percentage. Lipids with net charge <-1 were added so as to the overall charge density the same as with the corresponding MLVs with (−1) charged lipids. Chloroform was evaporated under a gentle stream of dry nitrogen gas. The dried lipid mixtures were left under vacuum overnight to completely remove all traces of chloroform and then hydrated with buffer (20 mM Tris-HCl pH 7.4, and 0.1 mM THP) to a final lipid concentration of 2 mg/ml.

### Giant unilamellar vesicles (GUVs)

GUVs were prepared as described previously (50). Briefly, a lipid mix was spread on the conductive side of the indium-tin oxide (ITO)-coated glass and left under vacuum overnight to remove all traces of chloroform. Electroformation was then performed in 600 mM sucrose solution for 1 h at 45°C at 1V, 10 Hz. All lipid mixes included cholesterol at 20 mol%, Atto647N-DOPE at 0.1 mol% and POPC as bulk lipid. POPS was included at 20 mol% and PI(4,5)P_2_ at 5 mol% to give the same nominal charge density on the membranes. To prevent segregation of PI(4,5)P_2_ from other lipids, the lipid mix and ITO-coated glass slide were preheated to 60 °C prior to spreading the lipids on the slides, and the electroformation was in this case performed for 1 h at 60°C.

### MLVs pulldown assay

CHIKV nsP1 in SEC buffer was added to MLVs suspension in 1:1 volume ratio keeping the final lipid concentration in the mixtures at 1 mg/ml. The lipid to protein ratio was kept at 500:1. The mixture was incubated at room temperature for 30 minutes and then centrifuged at 15000 rpm for 30 minutes at 4°C. The supernatant was carefully removed, after which equal amounts supernatant and pellet were run on 10% SDS-PAGE. After destaining the Coomassie stained gel, image was acquired with a Chemidoc Imaging System (Bio-Rad) and the relative intensity of bands were quantified using ImageLab software (Bio-Rad). Each experiment was repeated three times. Relative pellet intensity was used to calculate the MLVs bound-protein fraction and mean ± S.D was plotted using Prism (Graph-Pad).

### Confocal imaging

In a Lab-Tek II chambered coverglass (Fisher Scientific), 150 μL of GUVs were mixed with 150 μL of isosmotic buffer (20 mM Tris-HCl, pH 7.4, 300 mM NaCl, 0.1 mM THP) containing proteins at concentrations stated in Results. The mix was gently stirred and incubated 10 min at room temperature before imaging. Images were acquired using a Nikon A1R series confocal microscope equipped with a GaAsP detector and a Plan-Apochromat 60x oil (N.A 1.40) DIC objective. The ATTO647–DOPE membrane marker, and the ATTO488–labeled nsP1 were excited with 633-, and 488-nm lasers, respectively. z-stacks of GUVs were acquired at positions selected without observing the fluorescence channels. Each stack consisting of 10 images, spaced at 1 μm. Three experiment series were performed on three separate occasions with different batches of GUVs. For each series, images were acquired from total three wells, and from each well GUVs were imaged from 10 different field views. In each set of z stack, the nsP1 binding was calculated as the fraction of GUVs having visible nsP1 fluorescence above background. Data from all the three experiment series were then plotted against the respective GUVs types using Prism (Graph-Pad).

### nsP1-nsP2-membrane co-pelletation assay

MLVs of the lipid compositions POPC (10 %) : Cholesterol (20 %) : POPS (70 %) were used. In this assay, we kept the nsP2 concentration fixed to 0.55 µm and nsP1 concentration was titrated from 0 µM to 2.75 µM i.e., 1:5 in molar ratio. The final lipid concentration was kept at 1 mg/ml. The assay was performed as described above for MLV pulldown assay. The resulting gel was then silver stained. Images were acquired with a Chemidoc Imaging System (Bio-Rad) and the relative intensity of bands were quantified using ImageLab software (Bio-Rad). Each experiment was repeated two times. The pellet intensity at each nsP1 concentration was normalized to the total nsP2 intensity and plotted (mean ± S.D) against the nsP1 concentration using Prism (Graph-Pad).

### Mass Photometry

Mass photometry (MP) measurement was performed on a Refeyn OneMP (Refeyn Ltd.). Microscope coverslips (24 mm × 50 mm; Paul Marienfeld GmbH) were cleaned by serial rinsing with Milli-Q water and HPLC-grade isopropanol (Fisher Scientific Ltd.), on which a CultureWell gasket (Grace Biolabs) was then placed. For each measurement, 16 μL of buffer was placed in the well for focusing, after which 4 μL of nsP1 protein was added and mixed. The final protein concentration was 5 nM. Movies were recorded for 60 s at 100 fps under standard settings. Before measuring the protein sample, a protein standard mixture was measured to obtain a standard molecular weight calibration file. Data was processed using DiscoverMP software (Refeyn Ltd.).

## Supplementary figures

**Figure S1:**
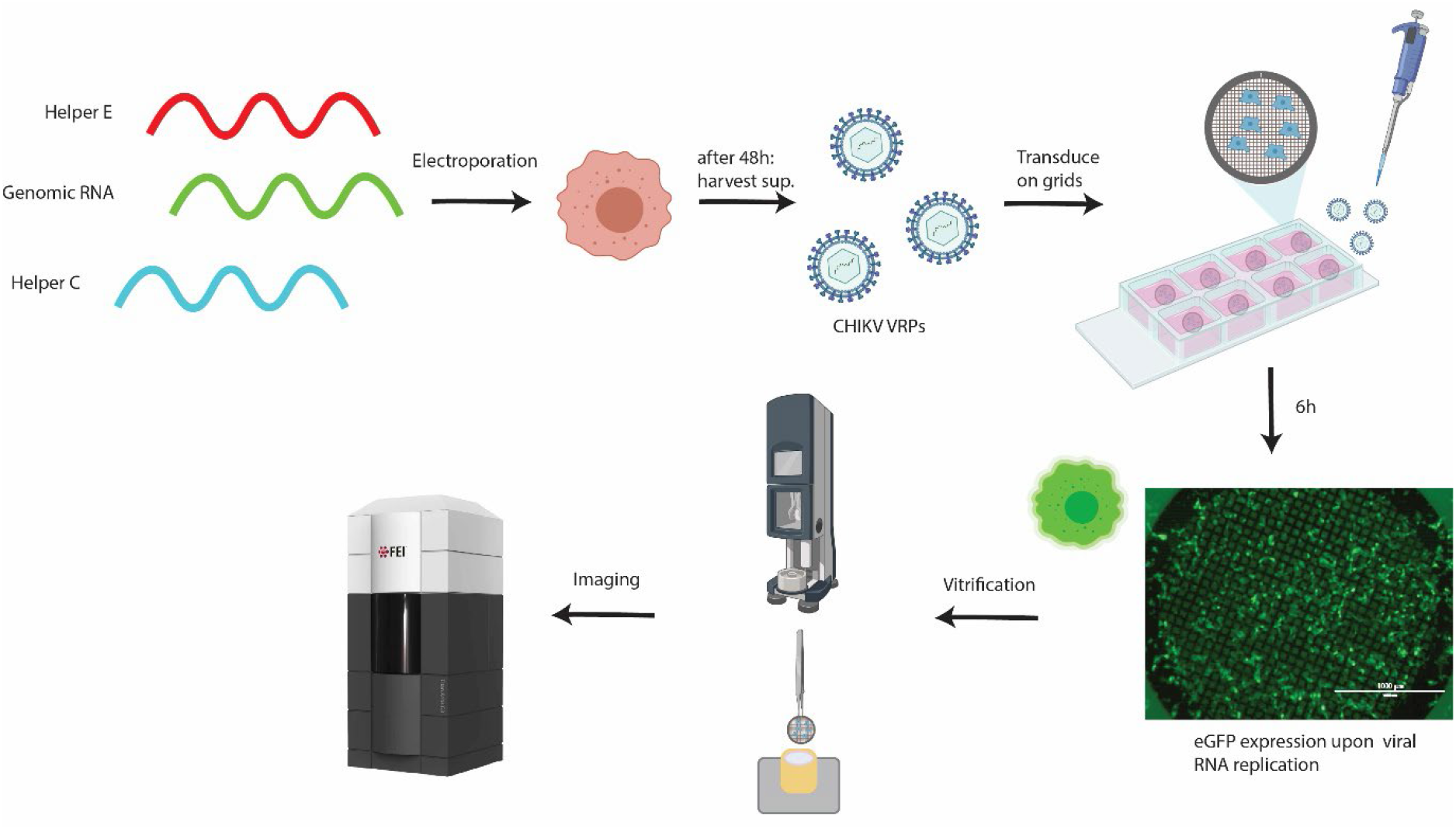
Workflow for cryo-electron tomography of CHIKV VRP-transduced cells. Production of CHIKV VRPs using three different RNAs (1). Green: genomic, carries the sequence of the non-structural proteins, and an eGFP sequence under regulation of the sub-genomic (SG) promoter. Red: “helper E”, carries the sequence of E1,E2,E3,6K/TF under regulation of the SG promoter. Blue: “helper C”, carries the sequence of the capsid protein under regulation of the SG promoter. See the Methods section for the detailed procedure. Cells electroporated with these three RNAs produce viral replicon particles (VRPs) that contain the only RNA having a packaging signal, i.e. the “genomic RNA” which is devoid of structural genes. BHK cells seeded on EM grids are transduced with CHIKV VRPs. 6h after transduction >∼90% of cells on grids display green fluorescence. After vitrification by plunge-freezing into liquid ethane-propane, cells are imaged at the Titan Krios cryo-TEM.

**Figure S2:**
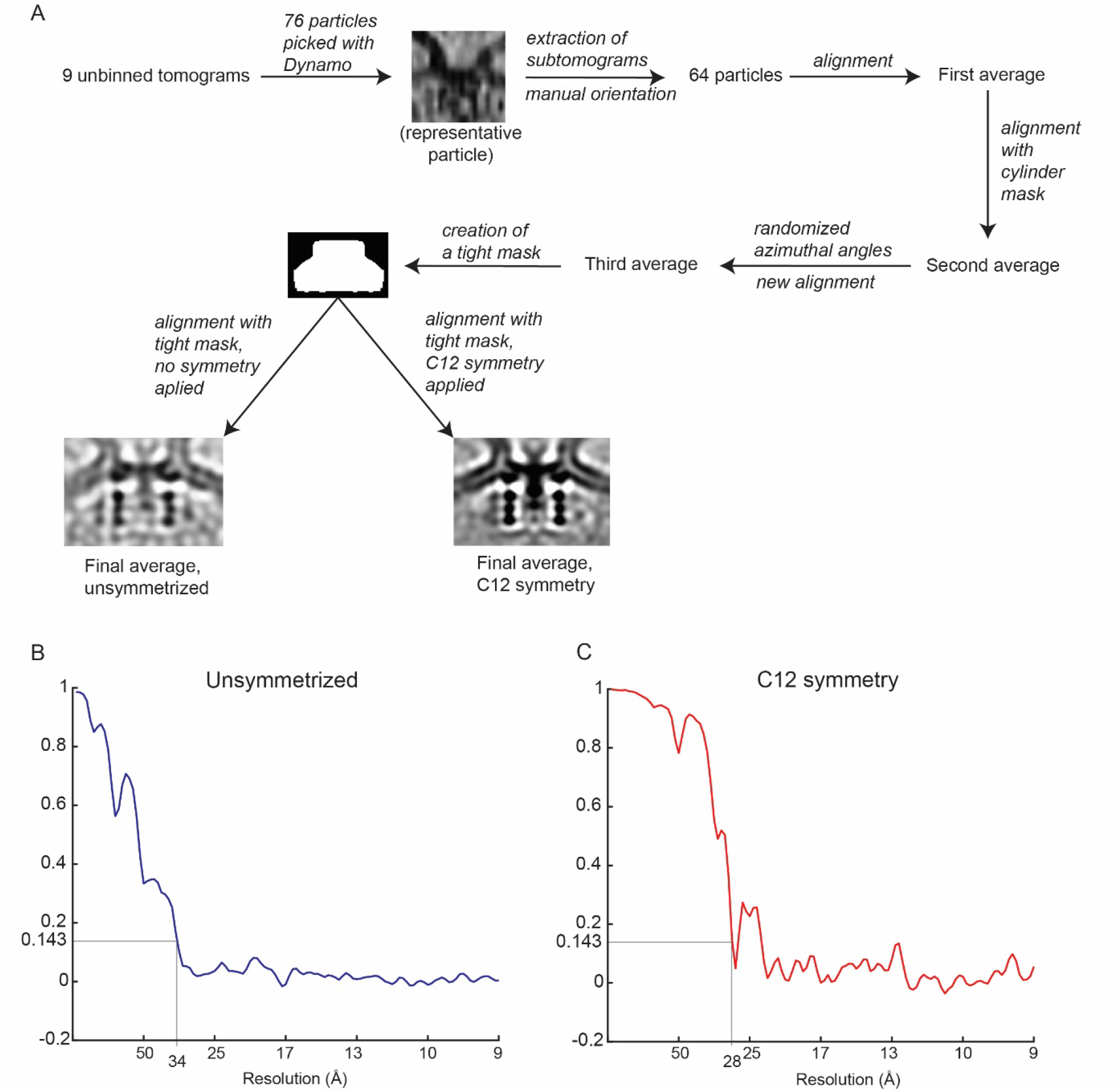
Subtomogram averaging of the neck complex. (A) Schematic of the subtomogram averaging process. (B) Gold-standard Fourier shell correlation of the unsymmetrized neck complex. (C) Gold-standard Fourier shell correlation of the symmetrized neck complex.

**Figure S3:**
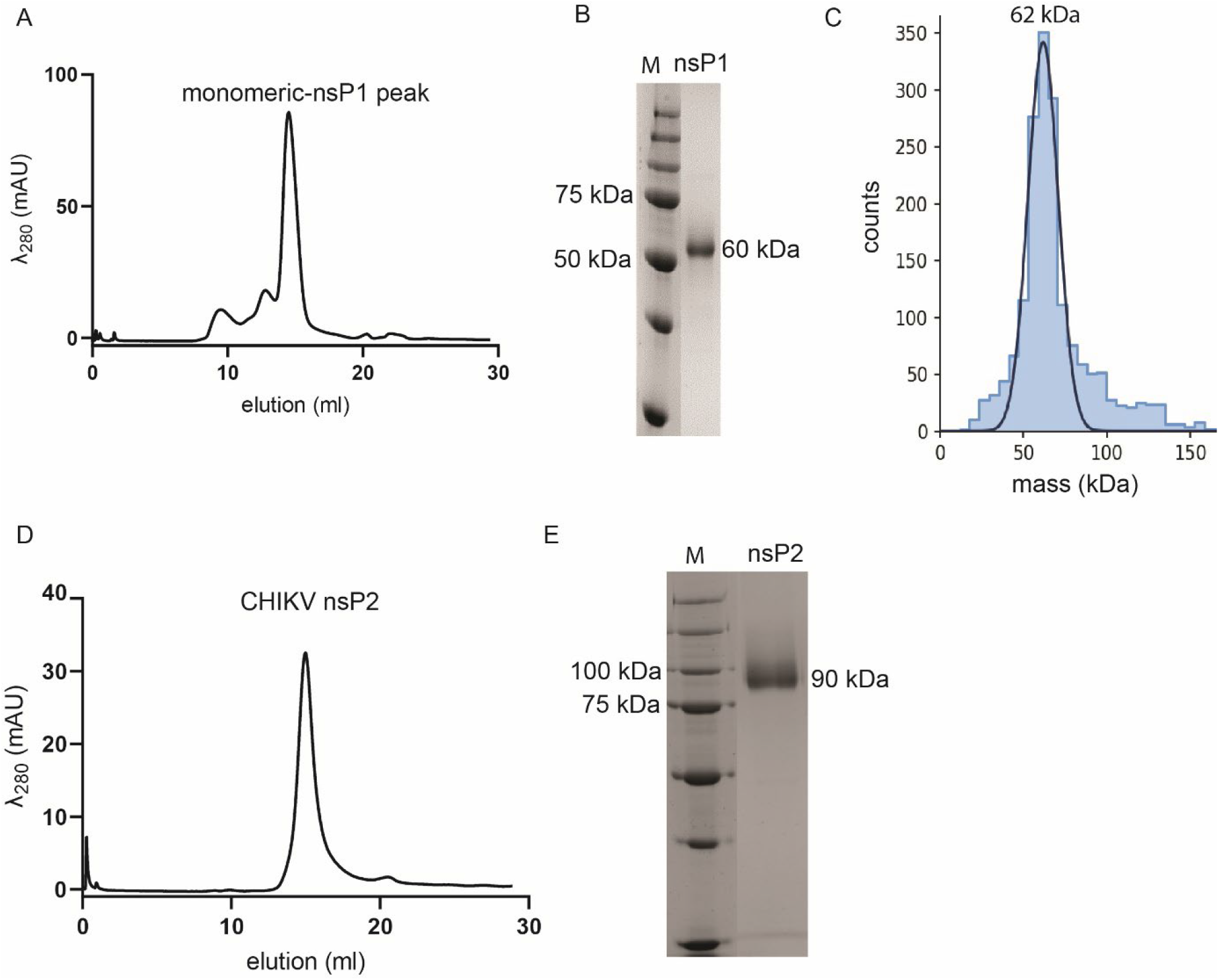
The purified CHIKV nsPs are homogenous and monomeric. (A) Size-exclusion chromatography (SEC) purification profile of CHIKV nsP1. The elution fraction at 14 ml represents the homogenous and monodisperse fraction of CHIKV nsP1, which was confirmed by Coomassie-stained 10 % SDS-PAGE (B) and mass photometry (C). Mass photometry shows that the protein has a molecular mass of 62 kDa, which is within the error range (± 5 %) from the expected mass of CHIKV nsP1 (60 kDa). (D) Size-exclusion chromatography (SEC) purification profile of CHIKV nsP2. The narrow elution peak represents the homogenous and monodisperse fraction of CHIKV nsP2, which was confirmed by Coomassie-stained 10 % SDS-PAGE (E).

**Figure S4:**
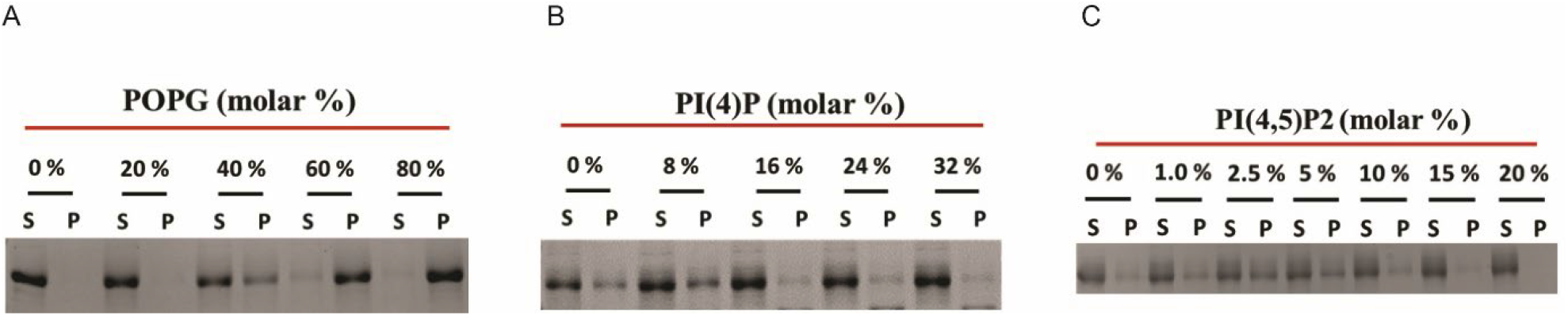
Representative gels related to Fig. 3. (A-C). Copelletation of nsP1 with multilamellar vesicles (MLVs) with varying percentages of the anionic phospholipids POPG (A), PI(4)P (B), and PI(4,5)P_2_ (C) in a background of POPC and 20% cholesterol. The supernatant and pellet were run on 10% SDS-PAGE. After destaining the Coomassie stained gel, image was acquired with a Chemidoc Imaging System (Bio-Rad) and the relative intensity of bands were quantified using ImageLab software (Bio-Rad) and plotted as shown in Figure 3 (A-C).

**Figure S5:**
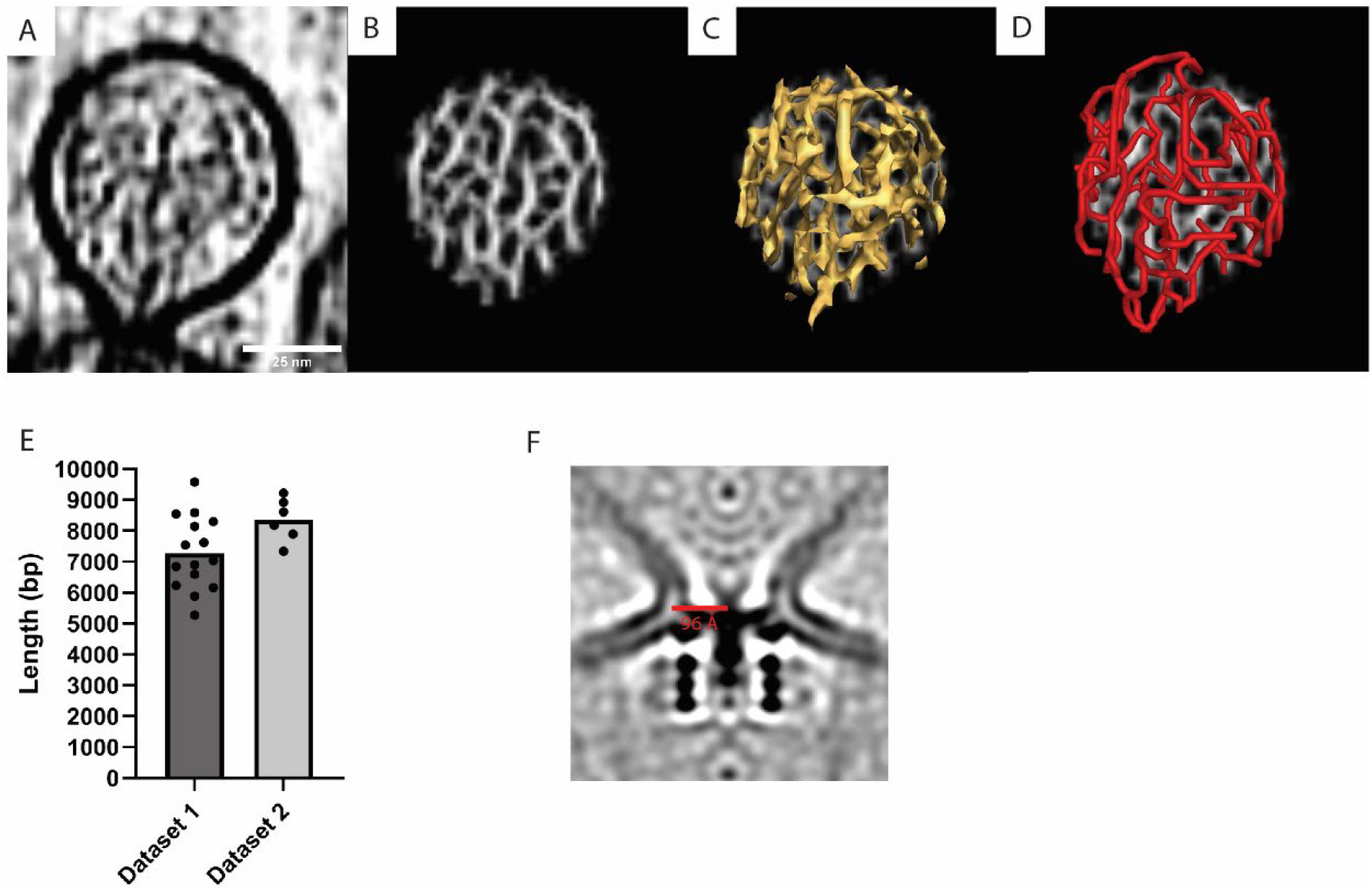
Tracing of RNA in the spherule lumen, and membrane neck diameter. (A) Slice through a tomogram highlighting the viral RNA in the spherule lumen. (B) The output of the cylinder correlation operation on the volume shown in (A), as performed in Amira’s filament tracing module (2). (C) Isosurface view of (B) (D) Filament model generated by correlation line tracing of the volume shown in (B). (E) Length of the traced RNA in Ångström. One dot corresponds to an individual spherule, and the bars represent the average value in each dataset (being tomograms acquired on different cells). (F) The radius of the neck of a spherule is 96 Å.

**Figure S6.**
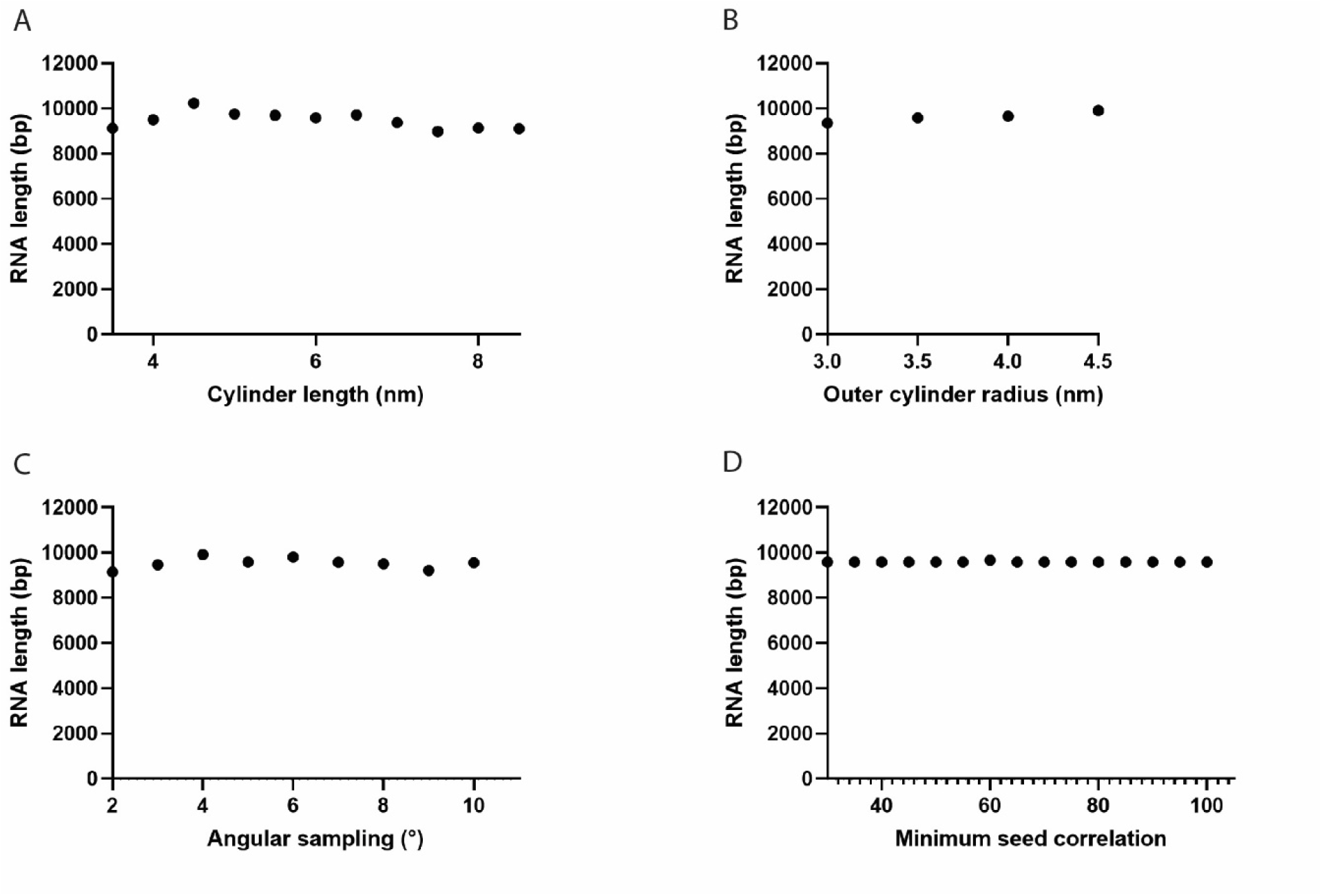
Dependence of the total filament length on filament tracing parameters. (A) Length of the cylinder templates used for cross-correlation caculation. A value of 6 nm was used in this study. (B) Radius of the outer cylinder mask. A value of 3.5 nm was used. (C) Maximum allowed angle between adjacent cylinder fragments during the tracing. A value of 5 degrees was used. (D) Cutoff value for the seed correlation between points following the RNA in cryo-tomograms. A value of 65 was used.

**Figure S7:**
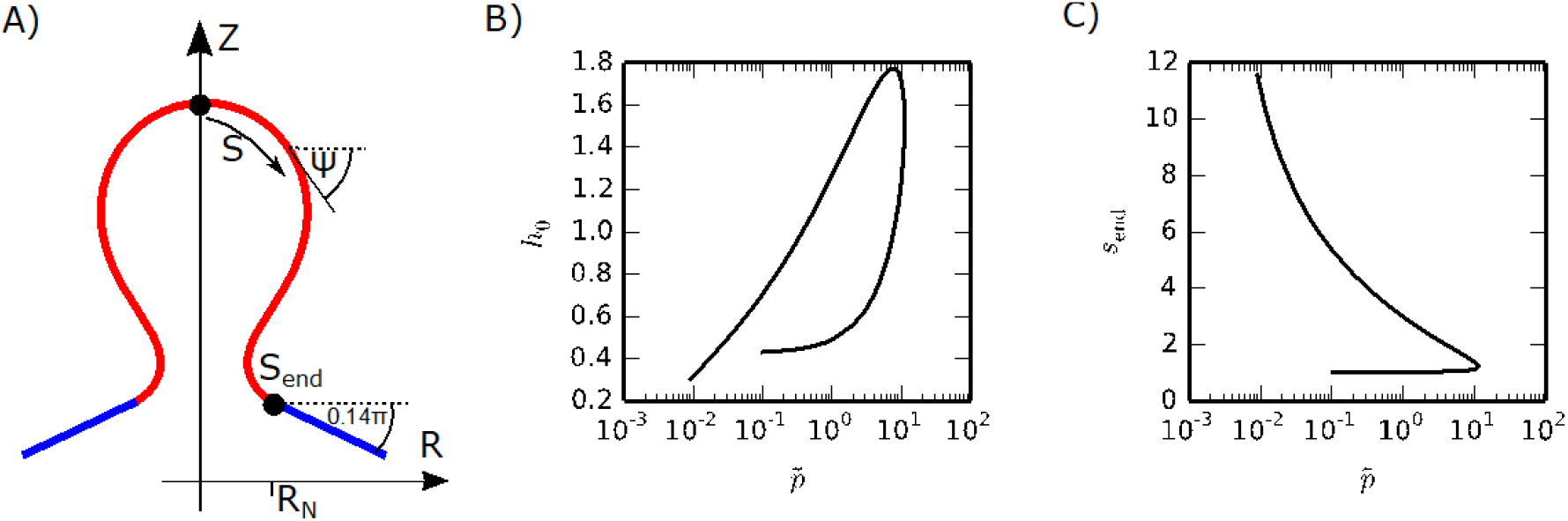
Spherule shape parametrization. (A) The spherule shape is parameterized by the arc length and the azimuthal angle ψ, where we consider a cylindrically symmetric shape. (B-C) values for *h*_0_ and *s*_*end*_ that solve the shape equations (Eqs. S17) with the boundary conditions, Eqs. (S19) for 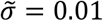.

