## Supporting Information for "Architecture of the chikungunya virus replication organelle"

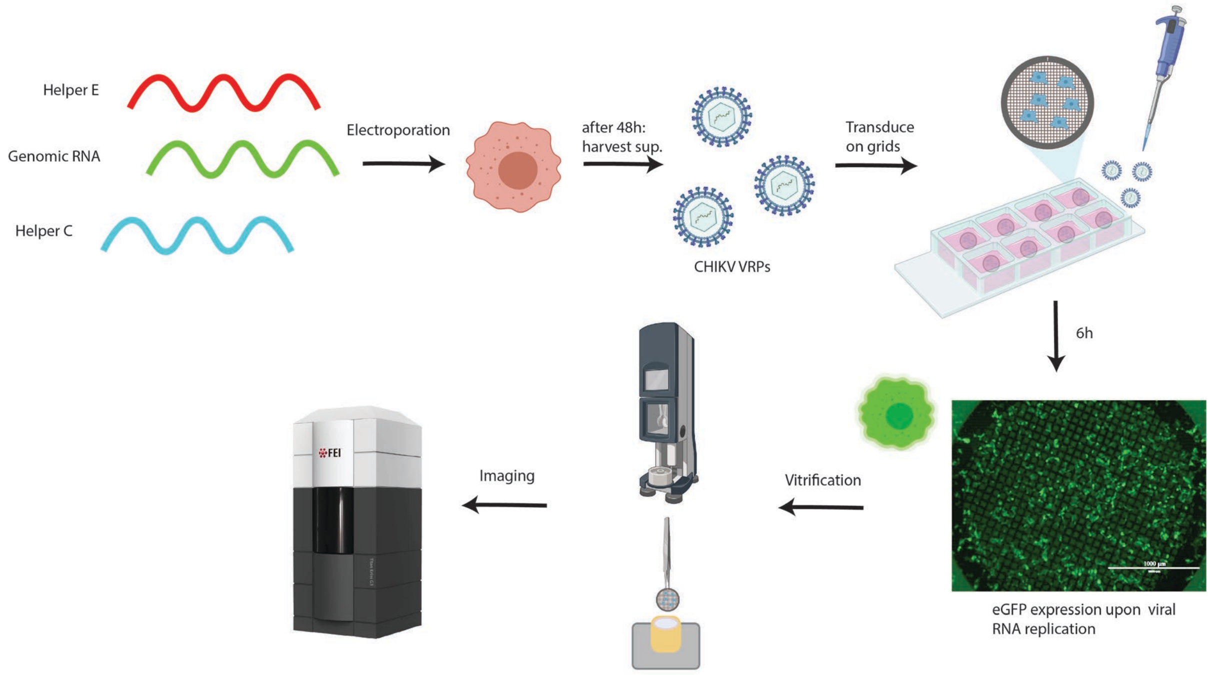


**Figure S1: Workflow for cryo-electron tomography of CHIKV VRP-transduced cells.** Production of CHIKV VRPs using three different RNAs (1). Green: genomic, carries the sequence of the non-structural proteins, and an eGFP sequence under regulation of the sub-genomic (SG) promoter. Red: “helper E”, carries the sequence of E1,E2,E3,6K/TF under regulation of the SG promoter. Blue: “helper C”, carries the sequence of the capsid protein under regulation of the SG promoter. See the Methods section for the detailed procedure. Cells electroporated with these three RNAs produce viral replicon particles (VRPs) that contain the only RNA having a packaging signal, i.e. the “genomic RNA” which is devoid of structural genes. BHK cells seeded on EM grids are transduced with CHIKV VRPs. 6h after transduction >~90% of cells on grids display green fluorescence. After vitrification by plunge-freezing into liquid ethane-propane, cells are imaged at the Titan Krios cryo-TEM.

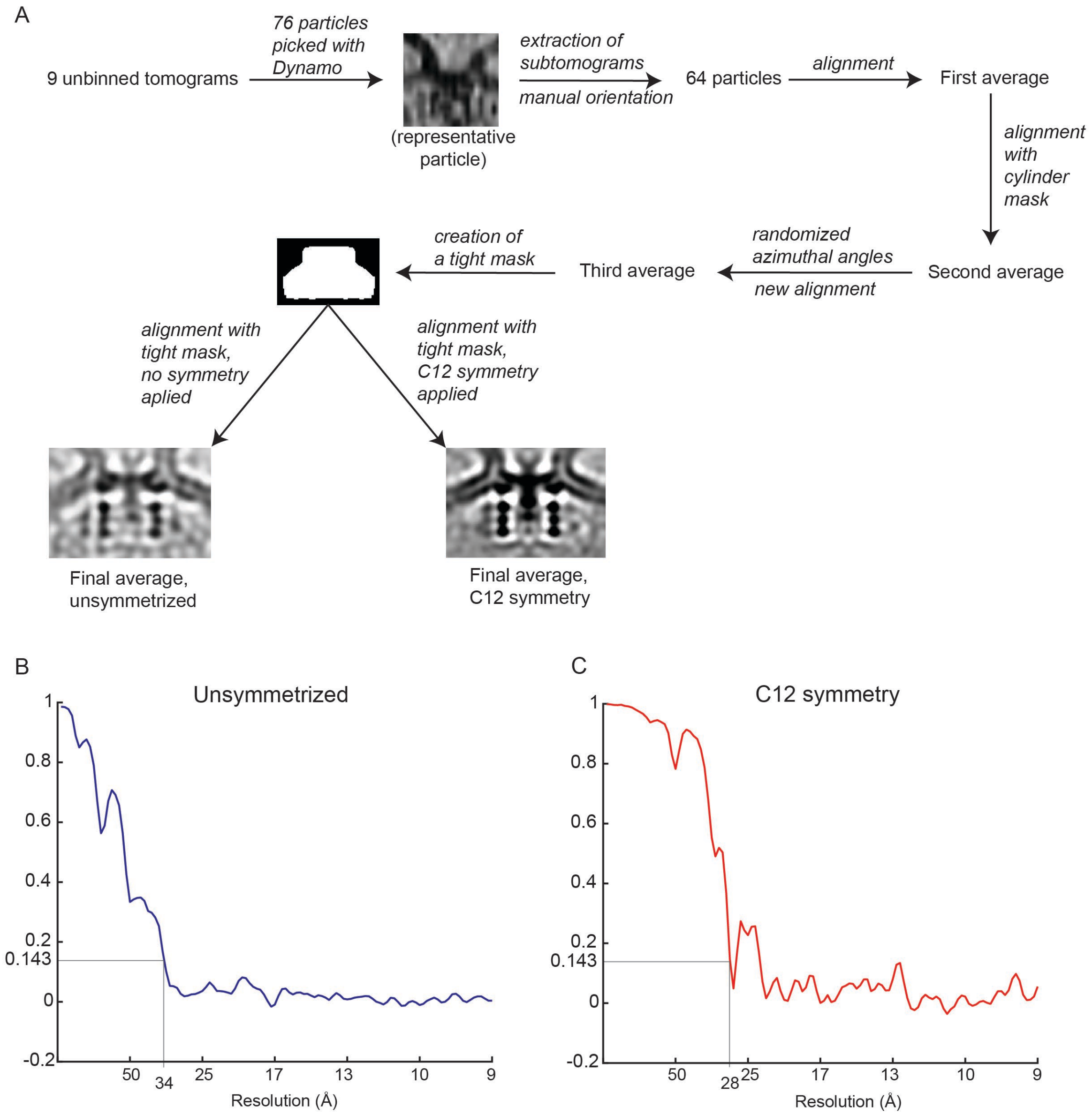


**Figure S2: Subtomogram averaging of the neck complex.** (A) Schematic of the subtomogram averaging process. (B) Gold-standard Fourier shell correlation of the unsymmetrized neck complex. (C) Gold-standard Fourier shell correlation of the symmetrized neck complex.

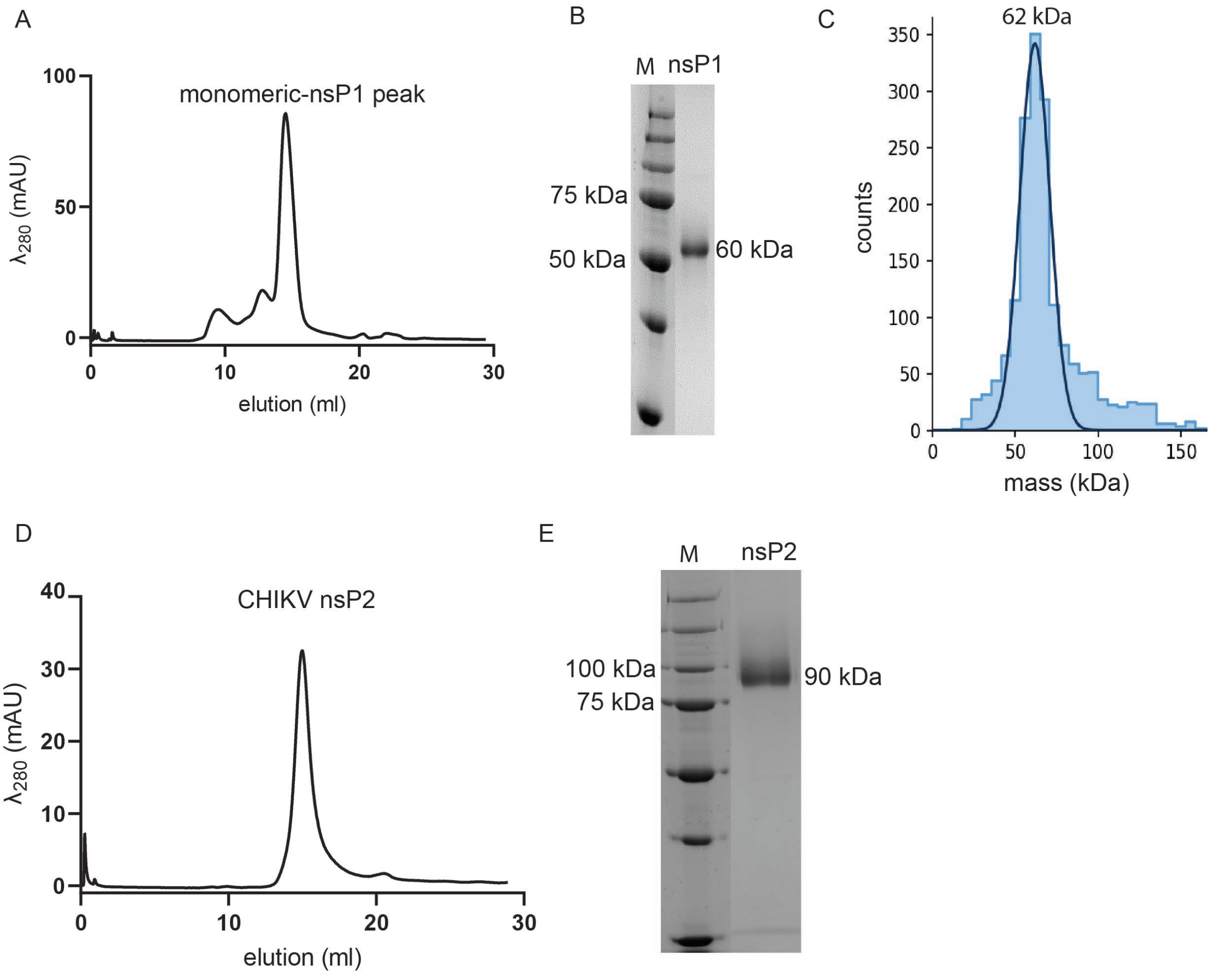


**Figure S3: The purified CHIKV nsPs are homogenous and monomeric.** (A) Size-exclusion chromatography (SEC) purification profile of CHIKV nsP1. The elution fraction at 14 ml represents the homogenous and monodisperse fraction of CHIKV nsP1, which was confirmed by Coomassie-stained 10 % SDS-PAGE (B) and mass photometry (C). Mass photometry shows that the protein has a molecular mass of 62 kDa, which is within the error range (± 5 %) from the expected mass of CHIKV nsP1 (60 kDa). (D) Size-exclusion chromatography (SEC) purification profile of CHIKV nsP2. The narrow elution peak represents the homogenous and monodisperse fraction of CHIKV nsP2, which was confirmed by Coomassie-stained 10 % SDS-PAGE (E).

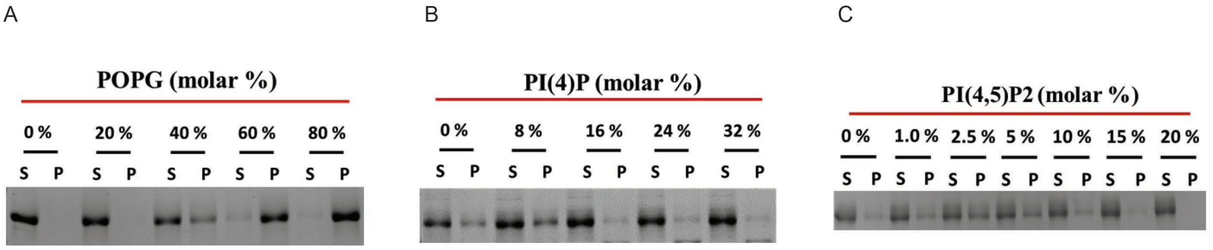


**Figure S4: Representative gels related to Fig. 3.** (A-C). Copelletation of nsP1 with multilamellar vesicles (MLVs) with varying percentages of the anionic phospholipids POPG (A), PI(4)P (B), and PI(4,5)$P_2$ (C) in a background of POPC and 20% cholesterol. The supernatant and pellet were run on 10% SDS-PAGE. After destaining the Coomassie stained gel, image was acquired with a Chemidoc Imaging System (Bio-Rad) and the relative intensity of bands were quantified using ImageLab software (Bio-Rad) and plotted as shown in Figure 3 (A-C).

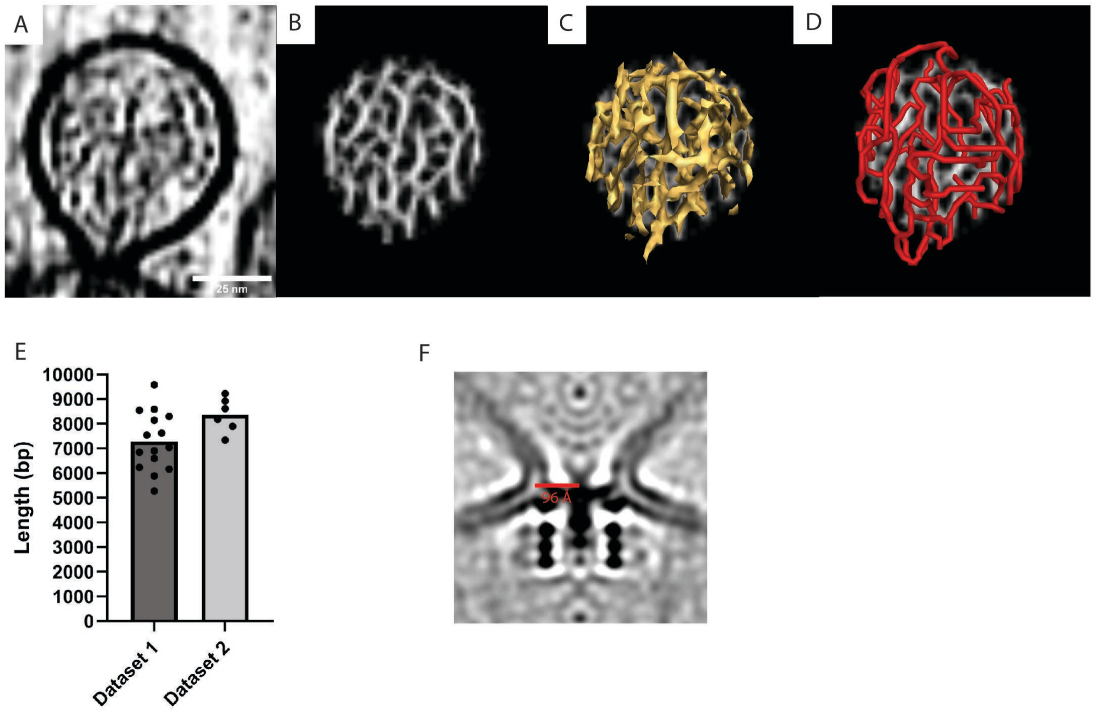


**Figure S5: Tracing of RNA in the spherule lumen, and membrane neck diameter.** (A) Slice through a tomogram highlighting the viral RNA in the spherule lumen. (B) The output of the cylinder correlation operation on the volume shown in (A), as performed in Amira's filament tracing module (2). (C) Isosurface view of (B) (D) Filament model generated by correlation line tracing of the volume shown in (B). (E) Length of the traced RNA in Ångström. One dot corresponds to an individual spherule, and the bars represent the average value in each dataset (being tomograms acquired on different cells). (F) The radius of the neck of a spherule is 96 Å.

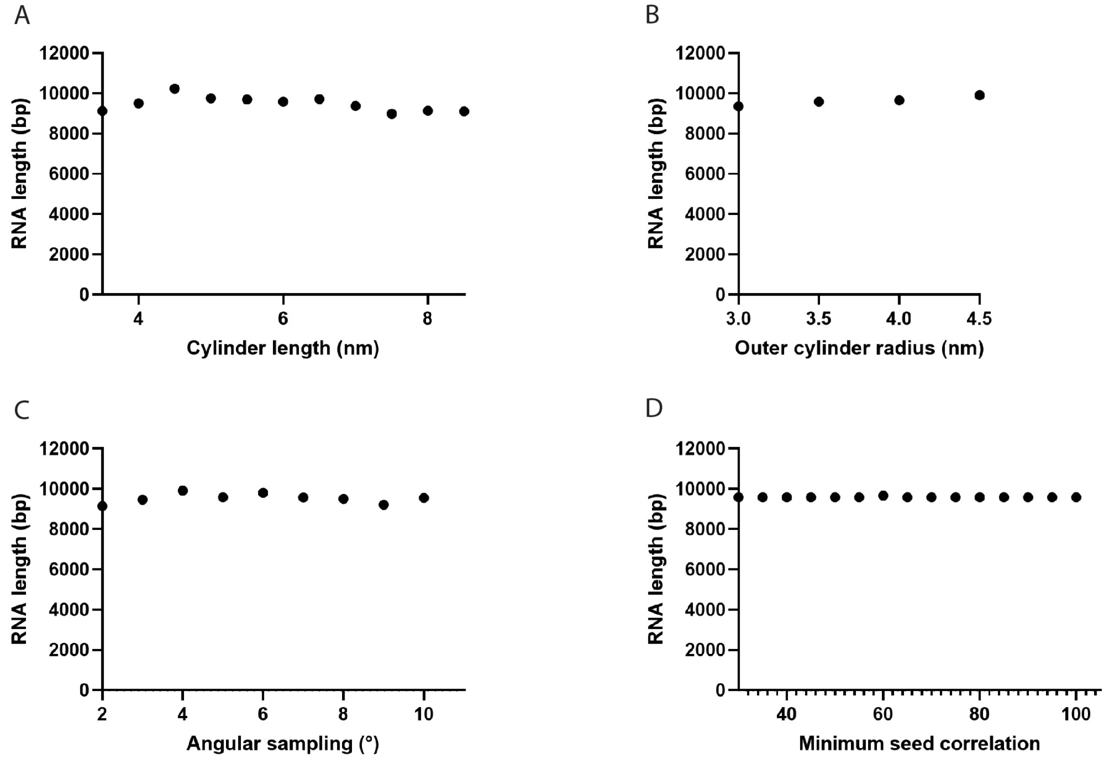


**Figure S6. Dependence of the total filament length on filament tracing parameters.** (A) Length of the cylinder templates used for cross-correlation caculation. A value of 6 nm was used in this study. (B) Radius of the outer cylinder mask. A value of 3.5 nm was used. (C) Maximum allowed angle between adjacent cylinder fragments during the tracing. A value of 5 degrees was used. (D) Cutoff value for the seed correlation between points following the RNA in cryo-tomograms. A value of 65 was used.

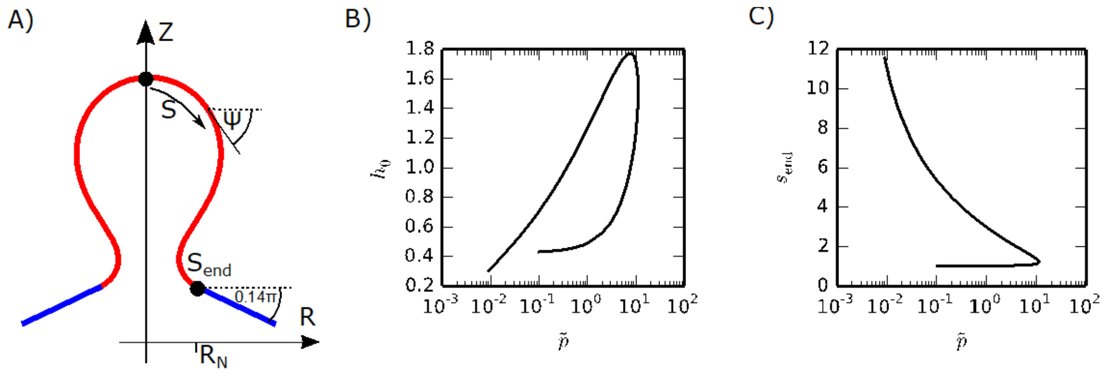


**Figure S7: Spherule shape parametrization.** (A) The spherule shape is parameterized by the arc length and the azimuthal angle $\psi$, where we consider a cylindrically symmetric shape. (B-C) values for $h_0$ and $s_{end}$ that solve the shape equations (Eqs. S17) with the boundary conditions, Eqs. (S19) for $\tilde{\sigma} = 0.01$.

**Table S1: Summary of the data collection parameters.**

| | |
|---|---|
| Data collection | |
| Microscope | Titan Krios G2 |
| Acceleration voltage (keV) | 300 |
| Camera | Gatan K2 Summit |
| Nominal magnification | 33,000 |
| Energy filter | Yes, BioQuantum |
| Slit width (eV) | 20 |
| Pixel size in super-resolution mode (Å) | 2.18 |
| Defocus range (µm) | -3 to -5 |
| Tilt range (°) | -60 to +60 |
| Tilt increment (°) | 2 |
| Total dose ($e^-/Å^2$) | 80 to 120 |
| Tomograms used for analyses | 9 |

**Table S2: Lipid compositions used to prepare multilamellar vesicles (MLVs).**

| **MLVs with (-1) charged lipids** | | | |
|---|---|---|---|
| **MLVs (equivalent charge %)** | **POPS/POPG/PI (mol %)** | **Cholesterol (mol %)** | **POPC (mol %)** |
| PS/PG/PI (0 %) | 0 | 20 | 80 |
| PS/PG/PI (20 %) | 20 | 20 | 60 |
| PS/PG/PI (40 %) | 40 | 20 | 40 |
| PS/PG/PI (60 %) | 60 | 20 | 20 |
| PS/PG/PI (80 %) | 80 | 20 | 0 |
| **MLVs with (-2.5) charged lipid** | | | |
| **MLVs (equivalent charge %)** | **PI(4)P (mol %)** | **Cholesterol (mol %)** | **POPC (mol %)** |
| PI(4)P (0 %) | 0 | 20 | 80 |
| PI(4)P (20 %) | 8 | 20 | 72 |
| PI(4)P (40 %) | 16 | 20 | 64 |
| PI(4)P (60 %) | 24 | 20 | 56 |
| PI(4)P (80 %) | 32 | 20 | 48 |
| **MLVs with (-4) charged lipid** | | | |
| **MLVs (equivalent charge %)** | **PI(4,5)$P_2$ (mol %)** | **Cholesterol (mol %)** | **POPC (mol %)** |
| PI(4,5)$P_2$ (0 %) | 0 | 20 | 80 |
| PI(4,5)$P_2$ (20 %) | 5 | 20 | 75 |
| PI(4,5)$P_2$ (40 %) | 10 | 20 | 70 |
| PI(4,5)$P_2$ (60 %) | 15 | 20 | 65 |
| PI(4,5)$P_2$ (80 %) | 20 | 20 | 60 |

**Predicting the relation between RNA length and spherule volume**

To derive a scaling relation between the length of the RNA and the volume of the spherule, we describe the spherule shape as a spherical cap (see inset in Fig.4 B in the main text), where the radius $R_s$ and the polar angle $\theta$ are related via the neck radius $R_N = R_s sin(\theta)$. The volume and area are then given by $V = \frac{\pi}{3} R_N^3 \frac{(2+\cos\theta)(1-\cos\theta)^2}{\sin\theta^3}$ and $A = 2\pi R_N^2 \frac{1-\cos\theta}{\sin\theta^2}$ and the membrane energy (Eq. (1) in the main text) reads

$$\frac{F}{\pi\kappa} = 4(1-x) + 2\frac{\sigma R_N^2}{\kappa}\frac{1-x}{1-x^2} - \frac{P R_N^3}{\kappa}\frac{(2+x)(1-x)^2}{3(1-x^2)^{\frac{3}{2}}},$$

( S1)

with $x = \cos\theta$. Minimization with respect to $x$ leads to

$$\frac{dF/(\pi\kappa)}{dx} = -4 - \frac{\sigma R_N^2}{\kappa}\frac{2}{(1+x)^2} + \frac{P R_N^3}{\kappa}\frac{1}{(1+x)^2\sqrt{1-x^2}} = 0$$

( S2)

and

$$\frac{P R_N^3}{\kappa} = 4(1+\cos\theta)^2 \sin\theta + 2\frac{\sigma R_N^2}{\kappa}\sin\theta.$$

( S3)

For a fully formed spherule, *i.e.* $\theta \approx \pi$, we write the pressure (Eq. S3) as a Taylor expansion around $\theta = \pi$:

$$\frac{P R_N^3}{\kappa} = (\pi-\theta)^5 + \frac{\sigma R_N^2}{\kappa}\left[2(\pi-\theta) - \frac{1}{3}(\pi-\theta)^3 + \frac{1}{60}(\pi-\theta)^5\right] + \mathcal{O}((\pi-\theta)^6)$$

( S4)

In analogy, the inverse of the spherule volume is expressed as a Taylor expansion around $\theta = \pi$:

$$\frac{R_N^3}{V} = \frac{3}{4\pi}(\pi-\theta)^3 + \mathcal{O}((\pi-\theta)^4)$$

( S5)

Inserting Eq. (S5) into Eq. (S4) we find:

$$P \approx \frac{\kappa}{R_N^3}\left(\frac{4\pi}{3}\right)^{\frac{5}{3}}\left(\frac{V}{R_N}\right)^{-\frac{5}{3}} + \frac{\sigma}{R_N}\left[2\left(\frac{4\pi}{3}\right)^{\frac{1}{3}}\left(\frac{V}{R_N}\right)^{-\frac{1}{3}} - \frac{1}{3}\left(\frac{4\pi}{3}\right)\left(\frac{V}{R_N}\right)^{-1} + \frac{1}{60}\left(\frac{4\pi}{3}\right)^{\frac{5}{3}}\left(\frac{V}{R_N}\right)^{-\frac{5}{3}}\right]$$

( S6)

Since $V \gg R_N^3$ for a mature spherule, the contribution to Eq. S6 that scale with the membrane tension are dominated by the $\left(V/_{R_N^3}\right)^{-1/3}$ term. Hence, Eq. (S6) simplifies to

$$P \approx \frac{\kappa}{R_N^3}\left(\frac{4\pi}{3}\right)^{\frac{5}{3}}\left(\frac{V}{R_N^3}\right)^{-\frac{5}{3}} + \frac{\sigma}{R_N} 2\left(\frac{4\pi}{3}\right)^{\frac{1}{3}}\left(\frac{V}{R_N^3}\right)^{-\frac{1}{3}}.$$

( S7)

From polymer theory it is known $P$, $V$ and $L$ the RNA length, scale as $PV{\sim}LV^{-2/3}$, or equivalently $L{\sim}PV^{5/3}$ (3-5). Inserting Eq. S5 and Eq. S7, we find

$$L{\sim}\frac{\kappa}{R_N^3}\left(\frac{4\pi}{3}\right)^{\frac{5}{3}}\left[1 + \frac{\sigma R_N^2}{\kappa} 2\left(\frac{4\pi}{3}\right)^{-\frac{4}{3}}\left(\frac{V}{R_N^3}\right)^{\frac{4}{3}}\right],$$

( S8)

which is equivalent to Eq. (2) in the main text

$$L = L_0\left[1 + \frac{\sigma R_N^2}{\kappa} 2\left(\frac{3}{4\pi}\right)^{4/3}\left(\frac{V}{R_N^3}\right)^{4/3}\right].$$

( S9)

**Membrane shape transformation**

To study the membrane shape transformation going from a flat membrane to a full-sized spherule, we derive the shape equations based on the Euler-Lagrange formalism. To this end, we describe the membrane shape in a cylindrically symmetric shape by an arc length parameterization (Fig. S7A) with the arc length $S$ and the azimuthal angle $\psi$. The height $Z$ and the radial coordinate $R$ are then obtained via $\frac{dR}{dS} = \cos\psi$, $\frac{dZ}{dS} = -\sin\psi$ and the principle curvatures are given by $C_1 = \frac{\sin\psi}{R}$, $C_2 = \frac{d\psi}{dS}$, with the mean curvature $H = (C_1 + C_2)/2$ (6). The membrane energy then reads

$$F = 2\pi \int_0^{S_{end}} dS \left[\frac{\kappa}{2} R \left(\frac{d\psi}{dS} + \frac{\sin\psi}{R}\right)^2 + \sigma R - P\frac{1}{2}R^2 \sin\psi\right]$$

( S10)

To determine the energy minimizing shape, we consider the functional $\tilde{F}$ with the Lagrangian-like function $\mathcal{L}$:

$$\tilde{F} = \int_0^{s_{end}} \mathcal{L}, \qquad \mathcal{L} = r\left(\psi' + \frac{\sin\psi}{R}\right)^2 + 2\tilde{\sigma}r - \tilde{p}r^2\sin\psi + \gamma(r' - \cos\psi) \tag{S11}$$

We used unitless variables $s = S/R_N$, $r = R/R_N$, $\tilde{\sigma} = \sigma R_N^2/\kappa$, $\tilde{p} = PR_N^3/\kappa$ and where derivatives with respect to $s$ are indicated as $\frac{d}{ds} = (\,)'$. Furthermore, we introduce the unitless variables $z = Z/R_N$ and $v = V/R_N^3$ which will be used further below. The Lagrange multiplier function $\gamma$ enforces the geometrical relation between $r$ and $\psi$. Using the Euler-Lagrange formalism (7, 8) we find based on $\frac{d}{ds}\frac{\partial\mathcal{L}}{\partial\psi'} = \frac{\partial\mathcal{L}}{\partial\psi}$

$$h' = \frac{\gamma}{4}\frac{\sin\psi}{r} - \frac{\tilde{p}}{4}r\cos\psi, \qquad \text{with } h = \frac{\psi' + \frac{\sin\psi}{r}}{2} \tag{S12}$$

And based on $\frac{d}{ds}\frac{\partial\mathcal{L}}{\partial R'} = \frac{\partial\mathcal{L}}{\partial R}$

$$\gamma' = 4h\left(h - \frac{\sin\psi}{r}\right) + 2\tilde{\sigma} - \tilde{p}r\sin\psi \tag{S13}$$

The spherule geometry requires the following boundary conditions

$$r(0) = 0, \qquad \psi(0) = 0, \qquad r(s_{end}) = 1, \qquad \psi(s_{end}) = 0.14\pi \tag{S14}$$

where the shape of the membrane neck is constrained by the protein complex to a radius $R_N$, i.e. $r(s_{end}) = 1$, and an angle $\psi = 0.14\pi$. To find a boundary condition for the Lagrange multiplier function $\gamma$ we determine the Hamiltonian-like function $\mathcal{H}$,

$$\mathcal{H} = -\mathcal{L} + \psi' \frac{\partial \mathcal{L}}{\partial \psi'} + r' \frac{\partial \mathcal{L}}{\partial r'} = rh\left(h - \frac{sin\psi}{r}\right) - 2\tilde{\sigma} r + \tilde{p} r^2 sin\psi + \gamma cos\psi \tag{S15}$$

We note that $\mathcal{H}$ is not an energy, but rather an auxiliary function that we use to derive an additional boundary condition. The explicit and implicit dependence of $\mathcal{H}$ and $\mathcal{L}$ on $s$ are related as $\frac{d\mathcal{H}}{ds} = -\frac{\partial \mathcal{L}}{\partial s}$. Since $\mathcal{L}$ does not depend explicitly on $s$, $\mathcal{H}$ is constant. The upper integration boundary $s_{end}$ is not fixed, which leads to $\mathcal{H} = 0$ (7, 8). From Eq. S15, we can now determine the boundary condition

$$\gamma(0) = 0 \tag{S16}$$

In summary, we obtain the following shape equations and boundary conditions

$$r' = \cos\psi \tag{S17a}$$

$$z' = -\sin\psi \tag{S17b}$$

$$\psi' = 2h - \frac{\sin\psi}{r} \tag{S17c}$$

$$\gamma' = 4h\left(h - \frac{\sin\psi}{r}\right) + 2\tilde{\sigma} - \tilde{p} r \sin\psi \tag{S17d}$$

$$h' = \frac{\gamma}{4}\frac{\sin\psi}{r} - \frac{\tilde{p}}{4} r \cos\psi \tag{S17e}$$

$$v' = \pi r^2 \sin\psi \tag{S17f}$$

$$r(0) = 0, \qquad z(0) = 0, \qquad \psi(0) = 0, \qquad \gamma(0) = 0, \qquad v(0) = 0$$

( S18a)

$$r(s_{end}) = 1, \qquad \psi(s_{end}) = 0.14\pi$$

( S18b)

Since Eq. (S16) has a singularity for $r = 0$, we shift the inner boundary from $s = 0$ to $s = \tau$. In the numerical calculations $\tau$ is set to $\tau = 0.001$. The mean curvature at the inner boundary is denoted as $h_0$. From $\psi(\tau) = \int_0^\tau \psi' \, ds \approx \int_0^\tau h_0 \, ds = h_0\tau$ we find the new boundary condition $\psi(\tau) = h_0\tau$. And from $r' = \cos\psi \approx 1 - \frac{\psi^2}{2} \approx 1 - \frac{(h_0\tau)^2}{2}$, we find $r(\tau) = \tau + \mathcal{O}(\tau^3)$. In analogy, we obtain the following boundary conditions at $s = \tau$ :

$$r(\tau) = 0, \qquad z(\tau) = 0, \qquad \psi(\tau) = h_0\tau\,, \qquad \gamma(\tau) = 0, \qquad h(\tau) = h_0, \qquad v(\tau) = 0$$

( S19a)

$$r(s_{end}) = 1, \qquad \psi(s_{end}) = 0.14\pi$$

( S19b)

For a given values of $\tilde{\sigma}$ and $\tilde{p}$ we have to find $h_0$ and $s_{end}$, such that the shape equations in Eqs. (S17) with the boundary conditions (Eqs. S19) are fulfilled. Values for $h_0$ and $s_{end}$ as a function of $\tilde{p}$ for $\tilde{\sigma} = 0.01$ are shown in Fig. S7B-C.
